# Jointly inferring the dynamics of population size and sampling intensity from molecular sequences

**DOI:** 10.1101/686378

**Authors:** KV Parag, L du Plessis, OG Pybus

**Affiliations:** Department of Zoology, University of Oxford, Oxford, OX1 3SY, UK; MRC Centre for Global Infectious Disease Analysis, Imperial College London, London, W2 1PG, UK

**Keywords:** coalescent model, sampling models, skyline plots, demographic inference, influenza, bison, Bayesian phylogenetics

## Abstract

Estimating past population dynamics from molecular sequences that have been sampled longitudinally through time is an important problem in infectious disease epidemiology, molecular ecology and macroevolution. Popular solutions, such as the skyline and skygrid methods, infer past effective population sizes from the coalescent event times of phylogenies reconstructed from sampled sequences, but assume that sequence sampling times are uninformative about population size changes. Recent work has started to question this assumption by exploring how sampling time information can aid coalescent inference. Here we develop, investigate, and implement a new skyline method, termed the epoch sampling skyline plot (ESP), to jointly estimate the dynamics of population size and sampling rate through time. The ESP is inspired by real-world data collection practices and comprises a flexible model in which the sequence sampling rate is proportional to the population size within an epoch but can change discontinuously between epochs. We show that the ESP is accurate under several realistic sampling protocols and we prove analytically that it can at least double the best precision achievable by standard approaches. We generalise the ESP to incorporate phylogenetic uncertainty in a new Bayesian package (BESP) in BEAST2. We re-examine two well-studied empirical datasets from virus epidemiology and molecular evolution and find that the BESP improves upon previous coalescent estimators and generates new, biologically-useful insights into the sampling protocols underpinning these datasets. Sequence sampling times provide a rich source of information for coalescent inference that will become increasingly important as sequence collection intensifies and becomes more formalised.

## I. Introduction

The coalescent process describes how the size of a population influences the genealogical relationships of individuals randomly sampled from that population (Kingman, 1982). Coalescent-based models are widely used in molecular epidemiology and ecology as null models of ancestry, and of the diversity of observed gene or genome sequences. In many instances, these sequences are sampled longitudinally through time from a study population, for example when individual infections are sampled across an epidemic caused by a rapidly-evolving virus or bacterium (Pybus and Rambaut, 2009), or when ancient DNA is extracted from preserved animal tissue that may be tens of thousands of years old (Shapiro and Hofreiter, 2014). If sequences accrue measurable amounts of genetic divergence between sampling times, then the dataset is termed heterochronous (Drummond *et al.*, 2003; Biek *et al.*, 2015). A common problem in molecular evolution is the estimation of effective population size history from these heterochronous sequences or from time-scaled genealogies (trees) that are reconstructed from those sequences.

Several coalescent-based approaches have been developed to solve this problem, including the popular and prevalent skyline and skygrid families of inference methods (Pybus *et al.*, 2000; Strimmer and Pybus, 2001; Drummond *et al.*, 2005; Minin *et al.*, 2008; Gill *et al.*, 2012). These approaches, which originated with the classic skyline plot of Pybus *et al.* (2000), estimate population size history as a piecewise-constant function using only the coalescent event times (i.e. the tree branching times) of the reconstructed genealogy. For heterochronous datasets, these methods typically assume that the sequence sampling times (i.e. the tree tips) are defined by extrinsic factors such as sample availability or operational capacity (Ho and Shapiro, 2011), and are thus uninformative about, and independent of, population size (Drummond *et al.*, 2005; Parag and Pybus, 2019).

Recent work has started to challenge this assumption and assess its consequences. Volz and Frost (2014) showed, for a coalescent process with exponentially growing population size, that including sequence sampling time information can notably improve the precision of demographic parameter estimates, if the sampling process is correctly specified. They recommended an augmented coalescent sequence-sampling model, and defined a *proportional sampling* process, in which the rate of sampling sequences at any time from a population is linearly dependent on its effective size at that time. Karcher *et al.* (2016) generalised this to include non-linear dependence, which they termed *preferential sampling*, and to allow for piecewise-constant effective population size changes. Karcher *et al.* (2016) cautioned that misleading inferences can result when the relationship between population size and the sequence sampling rate is misspecified.

While these works have brought attention to the benefits of exploiting sampling time information for population size inference, further progress is needed. Previous studies have treated the sampling model as a statistical addition to the coalescent process (Karcher *et al.*, 2016) and have not explicitly considered the types of sampling designs and surveillance protocols that are commonly implemented by epidemiologists and ecologists in the field. Moreover, there are to date few provable or general analytical insights into the joint inference of sampling and population size using coalescent models. A flexible model that can accurately assess the role of experimental and surveillance design is warranted as there is still uncertainty about what constitutes good rules for sequence sampling, and about the relative benefits and pitfalls of different sampling protocols (Stack *et al.*, 2010; Parag and Pybus, 2019; Hall *et al.*, 2016). These issues will only increase in importance as sequence sampling intensifies and heterochronous datasets become more common (Ho and Shapiro, 2011; Baele *et al.*, 2017).

Here, we aim to advance the field by developing a new sampling-aware coalescent skyline model, which we term the *epoch sampling skyline plot* (ESP). The ESP extends the classic skyline plot to include a flexible epoch-based sampling model that can represent biologically-realistic sampling scenarios. Its formulation also renders it amenable to theoretical exploration and straightforward implementation within a Bayesian phylogenetic MCMC framework. The ESP assumes that sampling occurs in epochs, which are defined as periods of time during which the sampling rate per capita is deemed constant. In practical applications, an epoch might, for example, represent weekly or monthly surveillance windows, epidemic seasons, archaeological periods or geological strata. The boundaries of each epoch are delineated by the sequence sampling times of the heterochronous genealogy. This guarantees model identifiability and helps guard against unsupported inferences (e.g. the number of per-capita sampling rate changes must be fewer than the count of sampling events).

Within an epoch, the ESP assumes that tree tips are sampled in proportion to population size, with a constant of proportionality that we call the sampling intensity. This intensity measures the average sampling effort over the epoch per capita, with larger values corresponding to faster rates of sequence sample accumulation. We allow the sampling intensity to change discontinuously between epochs, resulting in a flexible piecewise-constant sampling process. Within each epoch, the ESP locally models *density-defined sampling*, in which the sampling rate directly correlates with effective population size. Consequently, the ESP can describe a wide range of *time-varying*, density-defined sampling protocols. This allows the ESP to: (i) account for external, population-independent fluctuations in sampling effort and (ii) provide a means to quantify sampling effort through the testing of competing sequence-collection hypotheses. For example, we may be interested in whether sampling intensity increases or decreases through time, or whether known historical or experimental events are associated with a change in sampling intensity.

Although the flexibility of the ESP means it can model a wide range of sampling models, we here give attention to two specific sampling models, inspired by real-world collection practices. The first is density-defined sampling, which embodies the assumption that the availability of sequences depends on the size of the study population, and leads to a fixed proportion of the population being sampled across the sampling time frame of the study. It can be modelled in the ESP by forcing all epochs to have identical sampling intensities or, equivalently, by defining a single epoch that spans the entire sampling period. Density-defined sampling is a simple sequence-collection protocol and can be obtained directly from the proportional and preferential models of Volz and Frost (2014) and Karcher *et al.* (2016).

The second sampling model arises when studies aim to collect an approximately constant number of samples per unit time (e.g. week, epidemic season or geological era), irrespective of the size of the study population. This protocol is called *frequency-defined sampling* and is modelled within the ESP by allocating epochs uniformly over time, and allowing their individual sampling intensities to vary such that the process samples an (approximately) deterministic number of samples per epoch. Frequency-defined sampling is often undertaken in molecular epidemiology when resources for surveillance are limited or predefined; or when the primary research aim is to diagnose and classify infections or to provide snapshots of the genetic diversity of pathogen populations (Ho and Shapiro, 2011). Frequency-defined sampling cannot be described within previous frameworks and can represent the impact of external factors on the rate of sample collection.

In this paper we develop and define the ESP and show how it facilitates the joint inference of effective population sizes and sampling intensities, within maximum likelihood (ML) and Bayesian frameworks. We validate its performance using simulated data, before exploring its improvements over existing skyline-based methods on empirical datasets (H3N2 influenza A virus sequences from New York state, and ancient mtDNA sequences from Beringian steppe bison). We focus on biologically-inspired sampling protocols (see above) and demonstrate how the epoch-sampling model facilitates the testing and exploration of different data collection hypotheses. We highlight how the inverse relationship between the rates of sampling and coalescence can substantially improve population size estimation bias, especially when coalescent events are sparse, and prove that the information available for inferring population size (and hence the precision of those estimates) could more than double by using the ESP. Finally, we describe in detail both ML and Bayesian ESP implementations. The latter is available as an integrated package called BESP in the popular Bayesian phylogenetics platform BEAST2 (Bouckaert *et al.*, 2019).

## II. New Approaches

Consider a coalescent tree reconstructed from sequences sampled longitudinally through time. Let the effective population size underlying this process at time *t*, into the past, be *N*(*t*). Standard coalescent skyline-based approaches to estimating *N*(*t*) assume that sequence sample times are uninformative (Drummond *et al.*, 2005) and therefore draw all of their inferential power from the reconstructed coalescent event times. These methods approximate *N*(*t*) with a piecewise-constant function comprising *p* segments: 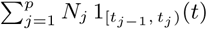, where *t*_*j*_ − *t*_*j−*1_ is the duration of the *j*^th^ segment and 1_𝔸_(*x*) is an indicator variable, which equals 1 if *x* ∈ 𝔸 and is 0 otherwise, for some set 𝔸. Here *t*_0_ = 0 is the present. Fig. 1 illustrates a coalescent sub-tree spanning the *j*^th^ segment, during which the effective population size is *N*_*j*_. Two epochs with distinct sampling intensities occur within this segment. The coalescent event times (grey) form the branching points of the reconstructed tree while sampling events (cyan) determine when new tips are introduced.

**Fig. 1:**
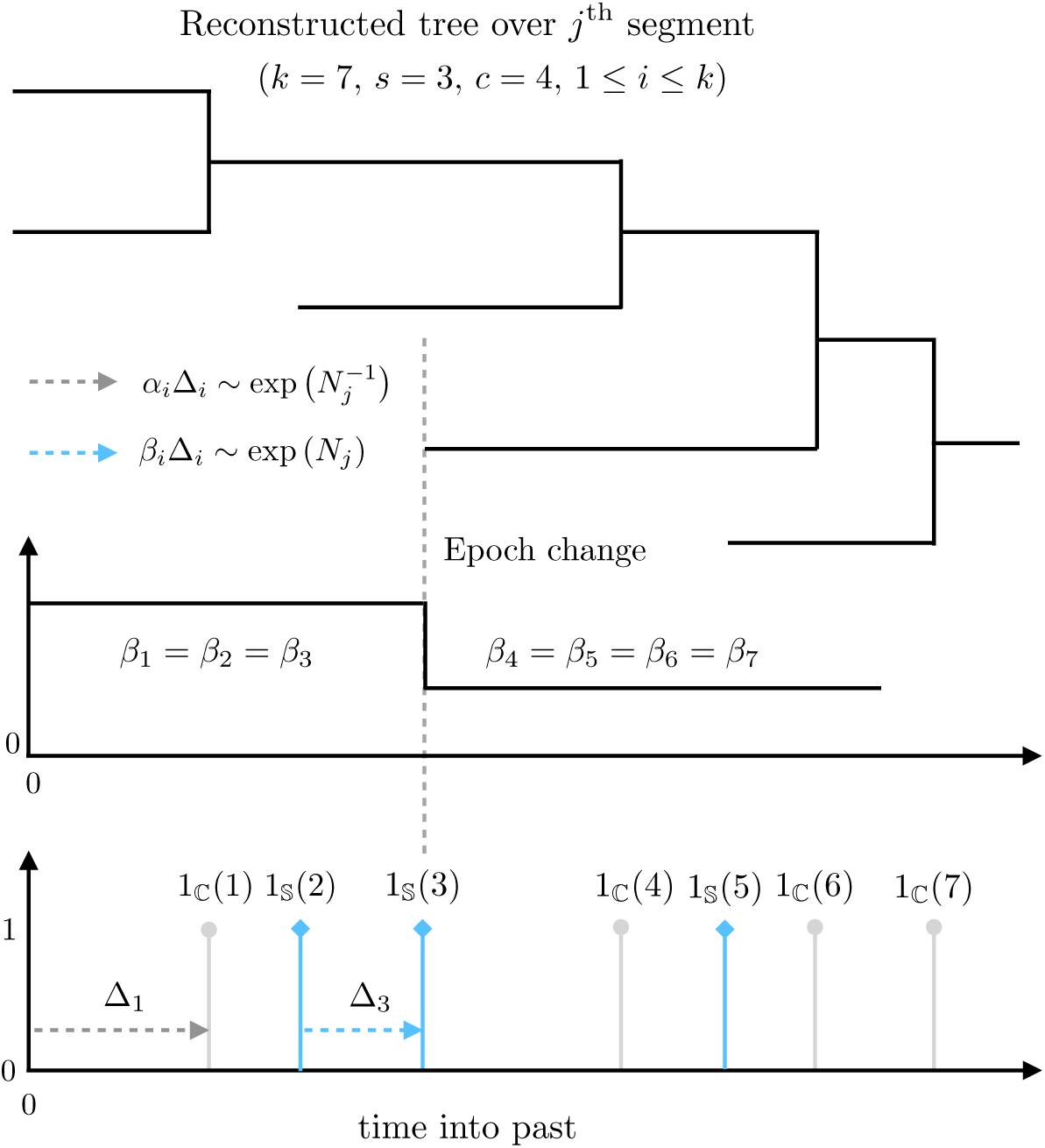
Illustration of the epoch sampling skyline plot model. A temporally sampled (heterochronous) tree consists of sampled tips and coalescing lineages. A portion of this tree is shown (top). This portion covers the *j*^th^ segment, during which the effective population size is assumed to be fixed at *N*_*j*_. Our epoch model assumes a piecewise-constant sampling intensity function which, in this illustration, comprises two epochs over this tree segment (middle). The sampling times (cyan, bottom) provide information about the sampling intensities in each epoch and also determine the epoch boundaries. The coalescent event times (grey, bottom) allow inference of *N*_*j*_ and also delimit the segment boundaries. See New Approaches for definitions of the mathematical notation used.

We use Δ_*i*_ to denote the duration of the *i*^th^ inter-event period or interval within a given segment, and define the lineage count in this interval as *ℓ*_*i*_. If there are *k* intervals in the *j*^th^ segment then 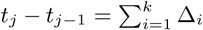. We use the sets 𝕊 and ℂ to indicate whether an interval ends with a sampling or coalescent event, respectively. Then 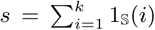 and 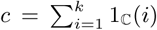 count the number of sampling and coalescent events in a given interval, and *k* = *s* + *c*. Note that *s, c* and *k* are not fixed, and can have different values for all *p* segments. Events which occur at a change-point belong to the interval that precedes that change-point i.e. the one closer to the present. Hence the sampling events 1_𝕊_(2) and 1_𝕊_(3) in Fig. 1 belong to the first epoch, and the starting two lineages are included in the likelihood of the (*j* − 1)^th^ segment.

Coalescent events falling within the *j*^th^ segment follow a Poisson process with rate 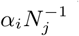, with 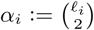, and *N*_*j*_, as the unknown effective population size during that segment (Kingman, 1982). As a result, 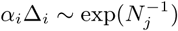 describes the key informative relationship in coalescent processes. Standard skyline methods capitalise on this dependence, but assume that intervals ending in sampling events (i.e. those satisfying {Δ_*i*_: *i* ∈ 𝕊}) are uninformative. Under this assumption the maximum Fisher information about *N*_*j*_ that can be extracted by these methods is 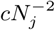 (Parag and Pybus, 2017).

The ESP instead posits that the sample times within the *i*^th^ interval of the *j*^th^ segment derive from a Poisson process of rate *β*_*i*_*N*_*j*_. Here *β*_*i*_ is the sampling intensity governing the average sampling effort (per capita or unit of *N*_*j*_) made across Δ_*i*_. This encodes the extra informative relationship: *β*_*i*_Δ_*i*_ ∼ exp(*N*_*j*_), and is the most complex sampling model admissible within the skyline framework (i.e. it is maximally parametrised). We remove unnecessary complexity by defining an epoch as a grouping of consecutive intervals (which may span multiple segment boundaries) over which the sampling intensity is constant. Thus, within an epoch, all *β*_*i*_ take the same value (in Fig. 1 there are two epochs). We force epoch change times to coincide with sequence sampling times, and assume that no sampling effort was made before the most ancient sample i.e. we set *β*_*i*_ = 0 for all intervals from the most ancient sample to the last coalescent event time (the time of the most recent common ancestor of the tree).

This description guarantees that the ESP is maximally flexible yet statistically identifiable (Parag and Pybus, 2019), since every skyline segment and epoch has at least one coalescent and one sampling event, respectively (see Results). Our epochal model, unlike previous attempts at incorporating sample times (Volz and Frost, 2014; Karcher *et al.*, 2016), can account for the temporal heterogeneity of sampling protocols undertaken in real-world studies. For example, sampling often occurs in bursts with discontinuous sampling effort that changes between collection periods. In the ESP, the sampling intensities of the epochs are independent of one other. Using this framework we construct the ESP log-likelihood for the *j*^th^ segment, ℒ_*j*_ = log P(𝒯 |*N*_*j*_), as in Eq. (1), with 𝒯 as the reconstructed tree.

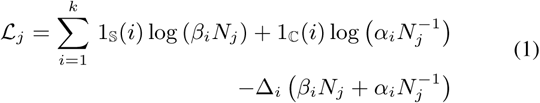

The complete tree log-likelihood is 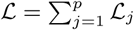. The waiting time until the end of any interval contributes the 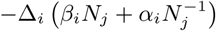 term, while sampling and coalescent events introduce terms 1_𝕊_(*i*) log(*β*_*i*_*N*_*j*_) and 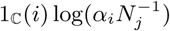, respectively. If we define *p*′ epochs over 𝒯, then there are *p* + *p*′ unknown parameters in our log-likelihood (the set of *N*_*j*_ and distinct, non-zero *β*_*i*_). Eq. (1) is related to the augmented log-likelihood from Karcher *et al.* (2016) but differs in both the population size and sampling models used.

The ESP is obtained from Eq. (1) by computing the grouped maximum likelihood estimate (MLE), 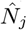, for each segment. This involves solving a pair of quadratic equations that depend on the relative number of sampling and coalescent events in that segment, *s* − *c*. Defining 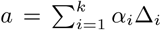 and 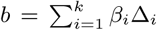, we obtain Eq. (2), from the roots of these quadratics (see Eq. (11) and Eq. (12) in Materials and Methods).

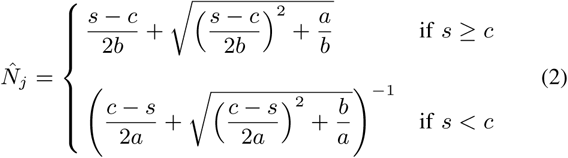

Eq. (2) forms our main result and requires the MLE of each 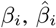, to be jointly estimated (see Materials and Methods for appropriate algorithms). If *s* = *c*, both parts of Eq. (2) converge to the simple square root estimator, 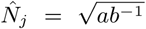. Grouping over *k* adjacent intervals in our skyline leads to smoother population size estimates that are quick to compute and easy to generalise. Note that if *s* = 0, all *β*_*i*_ = 0 and *c* = 1 then Eq. (2) simplifies to the classic skyline plot estimator of *N*_*j*_ (Pybus *et al.*, 2000).

The ESP has several desirable properties. Its counteracting proportional and inverse dependence on *N*(*t*) means that it has more informative intervals during time periods when coalescent events are infrequent, which otherwise hinders standard skyline inference. This property spreads the information about *N*(*t*) more uniformly through time, and reduces estimator bias. The ESP can also significantly improve overall estimate precision. The Fisher information that the ESP extracts from the *j*^th^ segment of the reconstructed tree is now at least 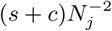 (see Results for analytic derivation).

## III. Results

### A. Simulated Performance

We start by comparing the estimates from Eq. (2) to those of the classic skyline plot (Pybus *et al.*, 2000), which ignores the information in sequence sampling times and is the basis of several popular skyline methods. We keep the number of piecewise-constant segments (parameters) inferred in the ESP (model dimensionality) approximately the same as that of the classic skyline plot by fixing *k* = 2. For clarity, we assume a single, known sampling intensity and examine only the period more recent than the most ancient sampling time. We compare the abilities of the ESP and classic skyline plot methods to recover a variety of population size dynamics in Fig. 2A–Fig. 2C. In each panel (A–C), the top graph gives the classic skyline plot estimate, the middle one shows the ESP estimate (for the same fixed tree), and the bottom one plots the distribution of sampling (cyan) and coalescent (grey) event times.

**Fig. 2:**
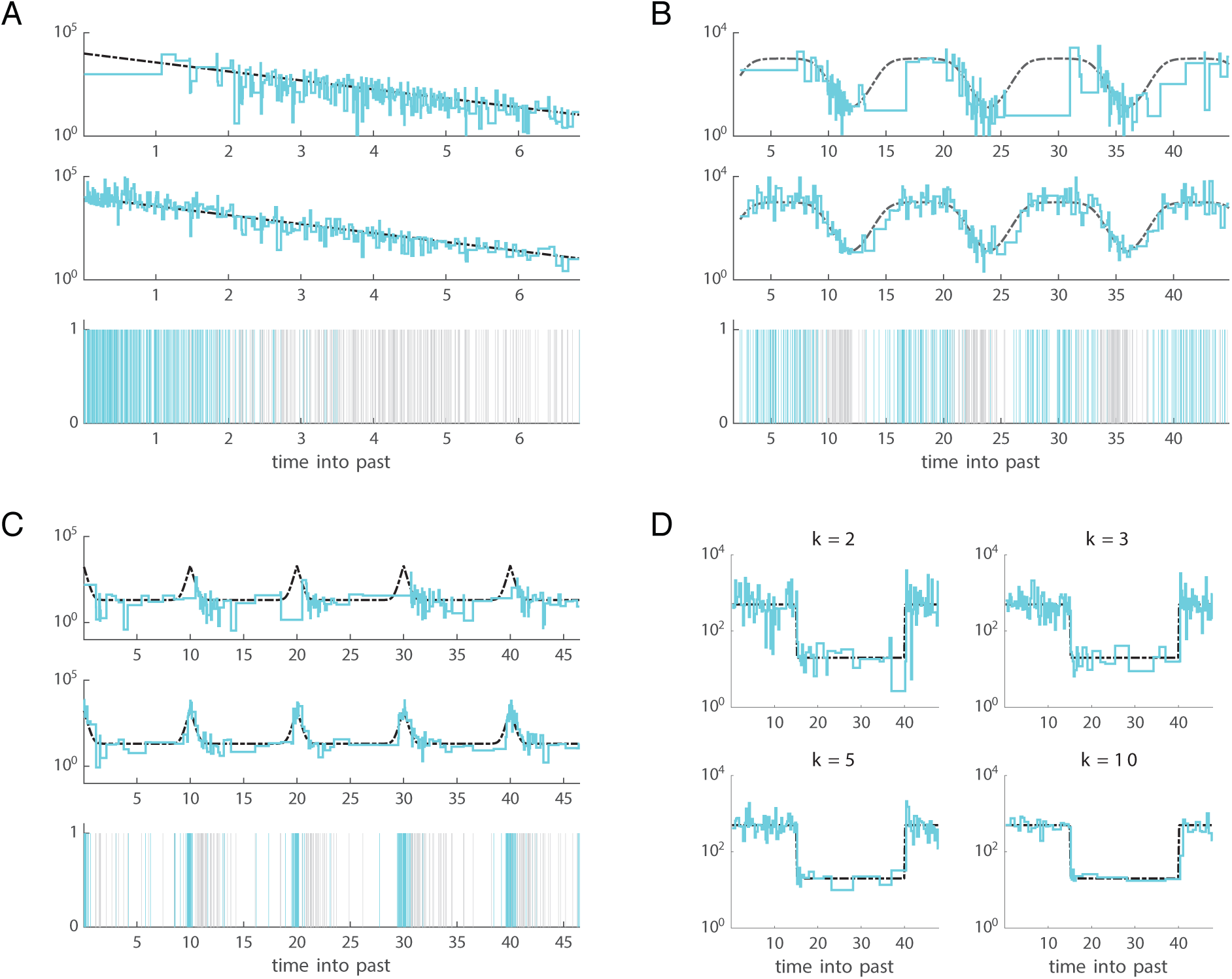
ESP and classic skyline plot estimates. Panels (A)–(C) compare the performance of the classic skyline plot (top graph) to the ESP at *k* = 2 (middle graph) for a range of demographic models: (A) exponential growth, (B) cyclical logistic growth and (C) steep periodic dynamics. Estimates of *N*(*t*) are shown in cyan and the true demographic functions are in dashed black, on a logarithmic scale. The classic skyline plot performs poorly near the present in (A), or when there are notable fluctuations between large and small population sizes in (B–C). This results from the uneven temporal distribution of coalescent events (grey lines in the bottom graph of each panel). The sampling events (cyan lines in the bottom graph of each panel) are inversely distributed to coalescent events. Consequently, the ESP tracks changes in population size more accurately, for the same number of population size segments. Panel (D) shows how increasing the grouping of adjacent intervals, *k*, can improve the smoothing of the ESP, in the context of a stepwise demographic function. All trees were simulated using the phylodyn R package (Karcher *et al.*, 2017) with approximately 300 coalescent and sampling events.

The ESP significantly improves demographic inference, relative to skyline plot methods, when population size is large (Fig. 2A) and in periods featuring sharp demographic changes (Fig. 2C). In these scenarios, standard skyline or skygrid approaches are known to perform poorly because coalescent events, due to their inverse dependence on population size, are sparse and hence unable to capture these population dynamics. Accordingly, coalescent events also tend to cluster around bottlenecks (Fig. 2B), and so cause standard methods to lose fidelity across cyclic epidemics. Sampling events, however, fall in periods of sparse coalescence, allowing the ESP to circumvent these problematic conditions.

The generalised skyline plot was introduced in Strimmer and Pybus (2001) to ameliorate the noisy nature of the classic skyline plot. It grouped adjacent intervals to achieve a bias-variance trade-off that led to smoother estimates of *N*(*t*). This grouping is used in some popular skyline approaches, notably the Bayesian skyline plot (BSP) (Drummond *et al.*, 2005). We achieve a similar smoothing effect in the ESP by increasing the grouping parameter, *k* (see Fig. 2D). This extends the generalised skyline plot approach in two ways; first by incorporating sampling time information, and second by including the specific times of events within a grouped interval.

Having clarified the attributes of the ESP, we now investigate examples in which the sampling intensities are unknown and can vary through time. We assume that the times corresponding to all sampling events are available for analysis. We consider two realistic, and widely-used sampling protocols, which we respectively refer to as *density-defined* and *frequency-defined* sampling. In the first there is a direct correlation between the time-varying effective population size and the rate of sampling, and a single sampling intensity persists throughout the complete sampling period. Density-defined sampling is the simplest model described within the ESP framework. It represents the process of proportional sampling (i.e. more samples are taken if the population to be sampled is larger).

However, in many epidemiological scenarios, surveillance organisations or treatment centres will often examine a relatively fixed number of samples per unit time (e.g. per month or epidemic season). This number may be constrained by extrinsic factors such as funding or operational capacity. Similar constraints may control the availability of ancient DNA sequences generated by molecular evolutionary studies. In such circumstances frequency-defined sampling results and the sampling intensity temporally fluctuates due to underlying changes in population size. Since this sampling scheme is more complex (it is a time-varying density-defined model), we use it to validate ESP performance. For clarity, in this section we restrict our analysis to fixed, time-scaled trees that are assumed to be known without error and apply our ML approach (see Materials and Methods for details). In later sections we examine both sampling models using a Bayesian implementation of the ESP that incorporates phylogenetic uncertainty.

We assume *p*′ epochs, so there are *p*′ unknown sets of *β*_*i*_ values to infer (within each epoch all *β*_*i*_ take the same value). We use *β* to represent this vector of unknowns, and let its MLE be 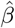. Note that epoch and population size change-points are not synchronised (i.e. they are generally non-overlapping), and we are jointly estimating a total of *p*+*p*′ parameters. Fig. 3A–Fig. 3D present our joint estimates of *N* and *β* at *k* = 20 for heterochronous genealogies simulated under four different demographic scenarios with frequency-defined sampling at *p*′ = 100 (Fig. 3A and Fig. 3C) or *p*′ = 50 (Fig. 3B and Fig. 3D) (see figure legend for details). Since the sample count in each epoch is approximately the same, the 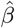 estimates (lower graphs of Fig. 3) take a complementary form to the 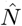 ones (upper graphs). These results show that the ESP has the ability to faithfully reproduce changes in both population size and sampling intensity through time.

**Fig. 3:**
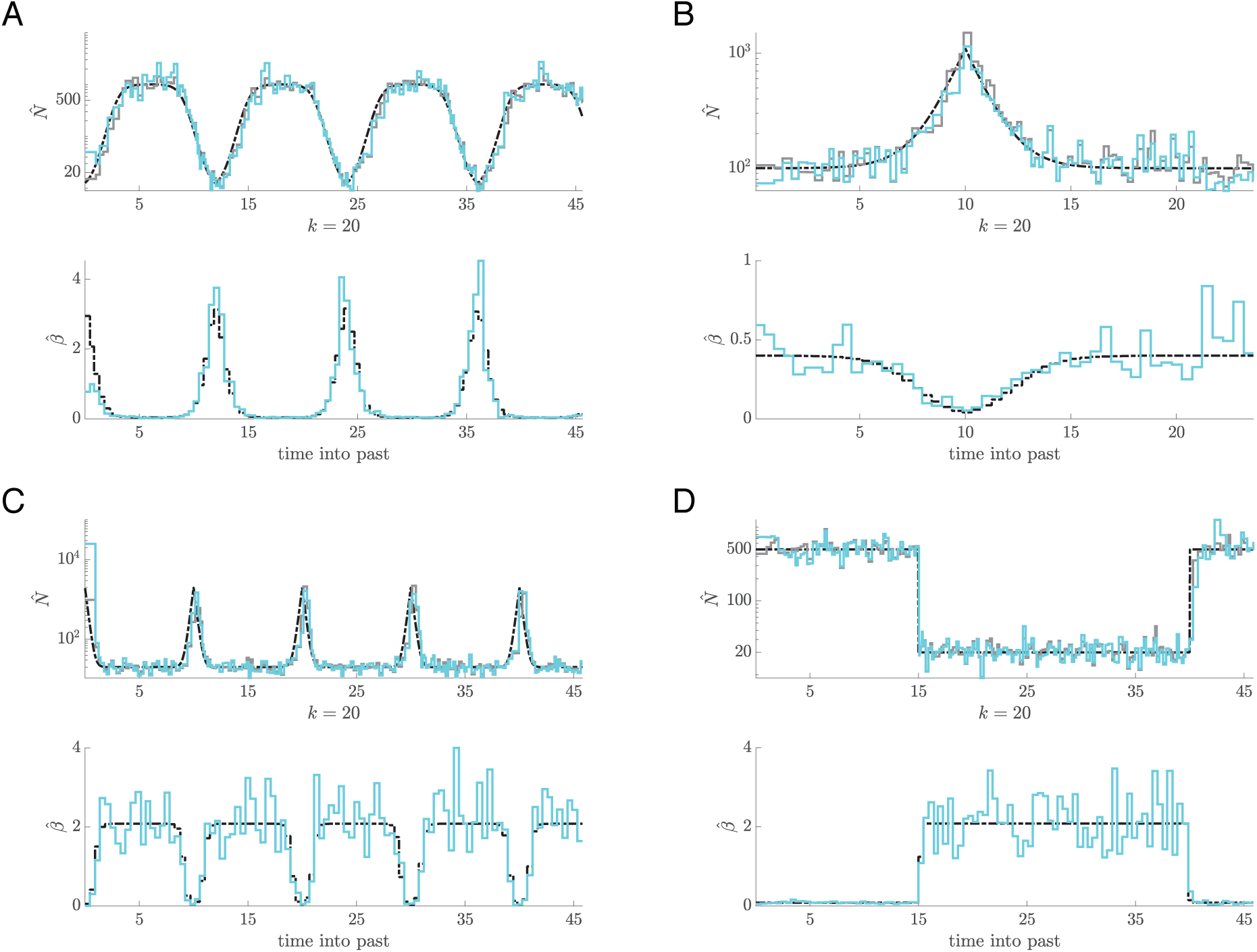
Joint inference of effective population size and sampling intensity using the ESP. Panels (A)–(D) show estimates of population size (upper) and sampling intensity (lower) through time. A single, fixed tree was simulated under a frequency-defined sampling model for demographic scenarios featuring: (A) cycles of logistic growth and decline (B) exponential growth and decline (the boom-bust model), (C) steep periodic cycles and (D) a step-wise population size change. Simulations in (A) and (C) comprised 2000 sampled tree tips over four population cycles, with *p*′ = 100 and *k* = 20. Simulations in (B) and (D) comprised 1000 sampled tree tips, with *p*′ = 50. The upper graph in each panel compares the true *N*(*t*) demographic function (dashed black) to 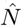 when *β* is known without error (grey), and 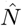 (cyan) when it is co-estimated with 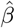 within the ML framework (see Eq. (2) and Materials and Methods). The lower graphs show the corresponding plots of 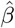 (cyan) against the true sampling intensity *β* (dashed black).

### B. Bayesian Implementation Simulation Study

Having explored the ML performance of the ESP, we now investigate and validate a Bayesian implementation of the ESP, which we call the BESP (see Materials and Methods). The BESP incorporates the ESP log-likelihood within the computational framework of BEAST2. In this section we benchmark the ability of the BESP to recover accurate and unbiased parameter estimates. We simulated 100 replicate coalescent genealogies (using the phylodyn R package (Karcher *et al.*, 2017)) under five demographic scenarios: (1) constant-size, (2) bottleneck, (3) boom-bust, (4) cyclical boom-bust and (5) logistic growth and decline. In all simulations we used frequency-defined sampling with approximately equal numbers of samples split over 24 equidistant epochs. We jointly inferred *N* and *β* from each simulated tree using the BESP and assumed that trees were known without error (to render the simulations computationally feasible, and to distinguish uncertainty in the coalescent model from phylogenetic noise). Estimation of *N* and *β* directly from sets of empirical gene sequences is demonstrated in the next section.

We grouped coalescent and sampling events into *p* = 100 equally-informed population size segments (i.e. *k* is equal for all segments) to estimate *N* and used *p*′ = 24 approximately equidistant sampling epochs for *β*. To quantify the bias and precision of the BESP method we computed the relative bias, the relative highest posterior density (HPD) interval width and the coverage of estimates of *N* and *β*, averaged across the time between the most recent and most ancient samples. Further details on the simulations, inferences and summary statistics can be found in the Supplementary Material. The results of our simulation study are summarised in Fig. 4. Example simulated trees and inferred parameter trajectories are shown in Figs. S1–S5^1^.

**Fig. 4:**
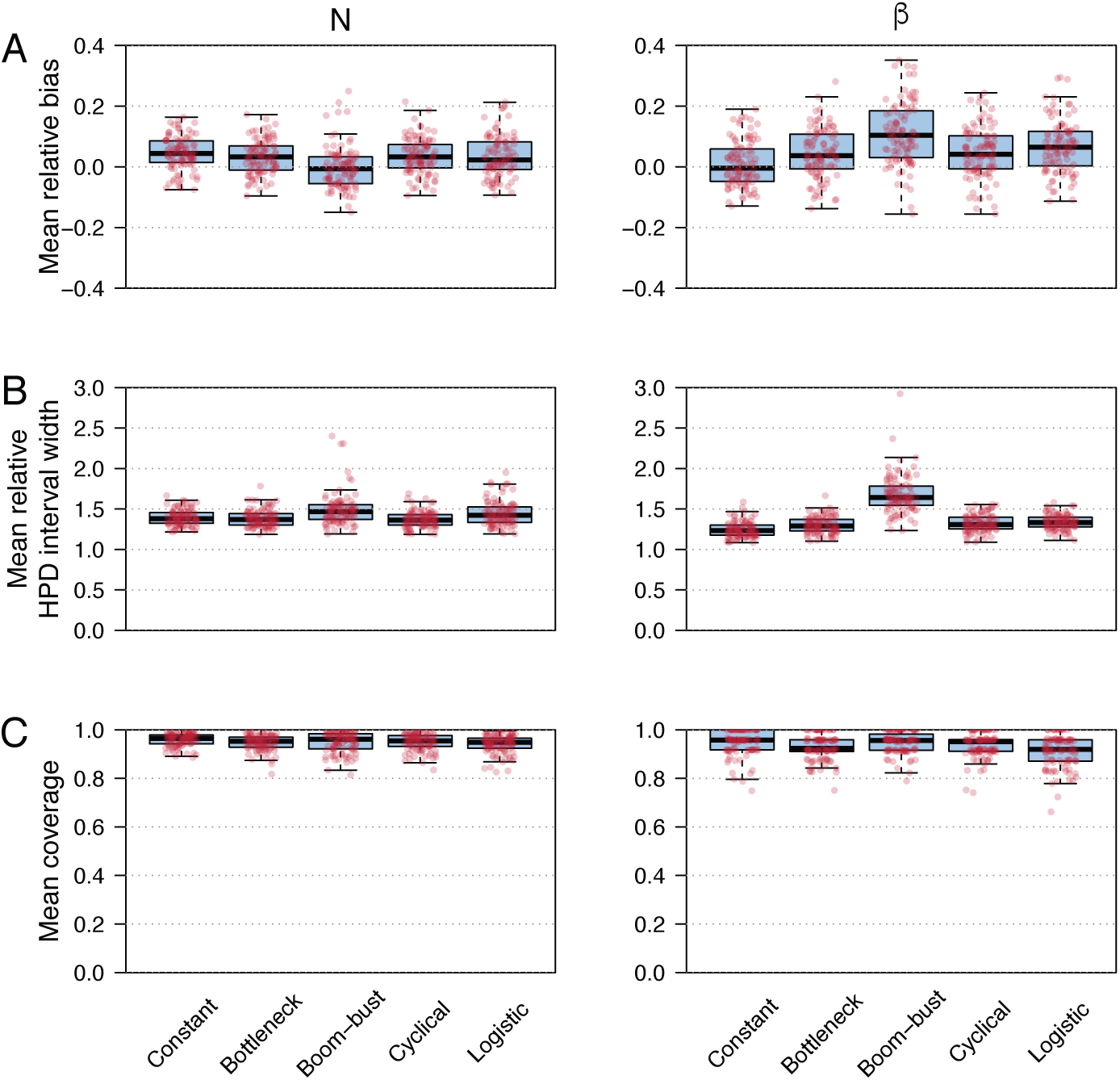
Boxplots and stripcharts showing measures of statistical performance for the BESP, evaluated on trees simulated under five different demographic models (constant, bottleneck, boom-bust, cyclical boom-bust, logistic growth and decline). We simulated 100 replicate trees for each scenario. Three measures of performance are shown (A) mean relative bias, (B) mean relative HPD interval size, and (C) mean coverage. Left and right columns illustrate estimation performance for effective population size (*N*) and sampling intensity (*β*), respectively.

Both *N* and *β* appear to be slightly overestimated with a larger bias in the *β* estimates. Nonetheless, the boxplots for the mean relative bias intersect 0 for all five demographic scenarios, verifying acceptable accuracy. The mean relative HPD interval widths of the population size estimates are < 2 for all replicate cases, with only a few outliers. Relative HPD intervals < 2 indicate that estimates are at least twice as precise as a standard Gaussian approximation (the width under a Gaussian distribution with standard deviation equal to the absolute value of the parameter is ≈3.92). Estimates of *β* under the boom-bust scenario occasionally have relative HPD interval widths > 2. We found this to be a consequence of the BESP not having sufficient power to precisely estimate *β* during the most recent sampling epoch (see Fig. S3 for an illustrative example of this effect). Lastly, the mean coverage is always close to 1, indicating that the true *N* and *β* values are included within the HPD intervals for the majority of the sampling period. These results verify that the BESP exhibits comparatively low bias and high precision.

### C. Case Study 1: Seasonal Human Influenza

Human influenza A virus (IAV) is a leading threat to global public heath, causing an estimated 290,000-650,000 deaths per year (WHO, 2018). Two subtypes of IAV currently co-circulate worldwide (H3N2 and H1N1-pdm) which, in temperate regions, cause annual winter epidemics. Strong immune pressure on the virus surface glycoprotein haemagglutinin (HA) drives a continuous replacement of circulating strains with new variants, termed antigenic drift (Ferguson *et al.*, 2003). Rambaut *et al.* (2008) reported 1,302 complete genomes of A/H3N2 and A/H1N1 viruses that were sampled longitudinally through time from temperate regions (specifically New York state, USA and New Zealand) and analysed the dynamics of IAV genetic diversity using the BSP (Drummond *et al.*, 2005).

Rambaut *et al.* (2008) found that the BSP could recover cyclical evolutionary dynamics from these sequences, with an increase in genetic diversity at the start of each winter influenza season, followed by a bottleneck at the end of that season, although the cycles were not sharply defined. Subsequently, Karcher *et al.* (2016) showed that estimates of IAV effective population size could be improved by incorporating sequence sampling time information within a preferential sampling model. However, that analysis assumed density-defined sampling and conditioned on the tree being known without error, thus eliminating phylogenetic noise. Here we extend the analysis of this dataset by using our BESP approach to co-estimate the effective population size history and sampling intensity across epidemic seasons of A/H3N2 HA genes sampled from New York state, USA.

As with the BSP, the population size parameter of the BESP, *N*, is proportional to the effective population size in the absence of natural selection (*N*_*e*_) i.e. *N* = *N*_*e*_*τ* where *τ* is the average generation time. This assumption does not hold for human IAV HA genes, which are subjected to strong directional selection. We follow previous practice and instead interpret *N* as a measure of relative genetic diversity (see e.g. Rambaut *et al.* (2008)). Our dataset comprises an alignment of 637 HA gene sequences (1,698 nt long) sampled across 12 complete influenza seasons, from 1993/1994 to 2004/2005 (Fig. 5A). Our estimates are inferred directly from the heterochronous sequence alignment using MCMC sampling and therefore incorporate phylogenetic uncertainty. Substitution and clock models are similar to those in Rambaut *et al.* (2008) (see Supplementary Material for model details). We estimate a BESP with *p* = 40 population size segments and *p*′ = 12 sampling epochs, so that each epoch corresponds approximately to the duration of one influenza season.

**Fig. 5:**
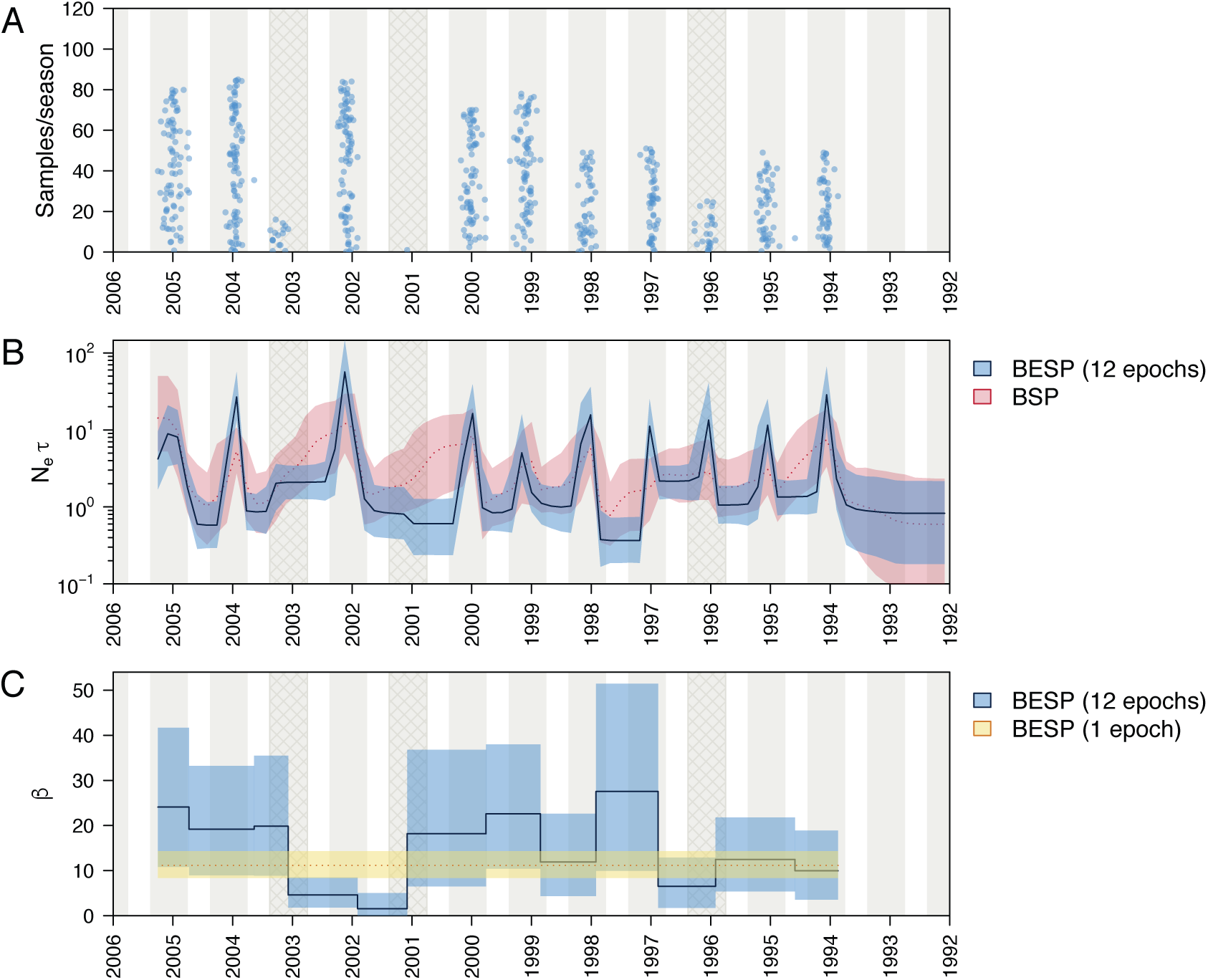
(A) Density of sequence sampling dates through time for the alignment of 637 A/H3N2 HA sequences from NY state that we analysed. Blue dots indicate stripcharts of individual samples for each season. The stripchart heights give the number of samples in each season. Grey shading indicates the approximate period of influenza observation in New York state during each season (epidemiological week 40, to week 20 in the next year). Cross-hatched seasons are those where A/H3N2 was not the dominant influenza virus subtype. (B) Median (solid/dotted line) and 95% highest posterior density (HPD) intervals (shaded areas) for the genetic diversity estimates (*N*_*e*_*τ*) through time. The BESP estimate is shown in blue and the BSP estimate is in red. (C) Median (solid line/dotted line) and 95% HPD intervals (shaded areas) of the estimated sampling intensities (*β*) for each sampling epoch. The 12-epoch BESP estimates are shown in blue and a single-epoch (density-defined) estimate is in yellow.

As Fig. 5A shows, considerably fewer sequences were sampled during the 1995/1996, 2000/2001 and 2002/2003 influenza seasons. The inferred dynamics of A/H3N2 genetic diversity (Fig. 5B) are strongly cyclical, with peaks coinciding with the midpoint of each epidemic season, except for 2000/2001 and 2002/2003. These results agree with epidemiological surveillance data for New York and New Jersey, which show that nearly all infections during the 2000/2001 season were caused by A/H1N1 and influenza B viruses, and that the 2002/2003 season was dominated by A/H1N1 infections (CDC, 2019). We do infer a clear peak for A/H3N2 in the 1995/1996 season (Fig. 5B), reflecting the fact that influenza cases during the 1995/1996 season were a mixture of A/H1N1 and A/H3N2 infections (Ferguson *et al.*, 2003), which resulted in an intermediate number of sequences being sampled that year (Fig. 5A).

A comparison of the BESP and BSP estimates of *N*_*e*_*τ* (Fig. 5B) on the same dataset shows that the BESP infers an epidemic peak for 1996/1997 whilst no such peak was revealed by the BSP. This indicates that the BESP has greater inferential power. Further, the peaks in the BESP are typically more defined than those in the BSP and have narrower 95% HPD intervals. Specifically, in the BESP, genetic diversity drops more sharply at the end of each season. This agrees well with our simulation results (see Fig. 2B), since coalescent events tend to be sparse when population sizes are large (e.g. at the start of a bottleneck), but sampling events are plentiful. Unlike the BESP, the BSP cannot exploit these informative sampling events and fails to efficiently track the fall in the number of infections.

The relative genetic diversity at the epidemic trough varies little among years, although it appears higher during 2002 and lower during 1997. It is possible that the bottleneck level largely depends on the availability of data since, in the absence of coalescent and sampling events, the smoothing prior maintains a roughly constant population size estimate (Volz and Frost, 2014). Since the informative events in a given season mostly stem from sequences sampled during that season (see Figs. S7 and S8), the BESP reveals no information about population dynamics prior to the first sampled season (1993/1994).

The inferred sampling intensities, *β*, for each season are given in Fig. 5C. Except for the period from 2001 to 2003 (which includes both of the seasons without an inferred epidemic peak), the 95% HPD intervals of the estimated *β* values for each season are overlapping. The estimated *β* for 1996 also appears lower, however the 95% HPD interval still overlaps with other seasons. Although there is some variation in the median estimates, the uncertainty in these estimates is large, especially when *β* is high.

We also analysed the same dataset using a simpler single-epoch model (i.e. density-defined sampling with a constant *β* through time). We found that the *N*_*e*_*τ* dynamics estimated using this simpler model (Fig. S6B) closely matches those inferred using the more complex 12-epoch model. The estimated sampling intensities obtained under the single-epoch and 12-epoch models are also congruent (Fig. 5C and S6). The density-defined model estimates a median sampling intensity of 11.16 (95% HPD 8.38–14.32), while the mean median estimate of the 12-epoch model is 14.87 (mean 95% HPD 6.03–27.19). We conclude that variation in sampling intensity through time is comparatively weak. Thus, if the aim of the original study authors was to undertake a density-defined sampling protocol then our *β* estimates provide an independent validation that this aim was, at least approximately, achieved.

### D. Case Study 2: Steppe Bison

To illustrate the application of the ESP model to non-virus datasets, we now analyse a heterochronous alignment of mtDNA genomes from modern and ancient bison that has previously been used to evaluate the performance of skyline-based methods (Shapiro *et al.*, 2004; Drummond *et al.*, 2005; Gill *et al.*, 2012; Faulkner *et al.*, 2019). During the Late Pleistocene, Beringia (eastern Siberia, the Bering land bridge, Alaska, and northwestern Canada) supported a large diversity of megafauna including bison, horses and mammoths. A favourable climate for specimen preservation means that bison fossils suitable for ancient DNA extraction are abundant across the region (Shapiro *et al.*, 2004). Sequences tens of thousand of years old can be recovered and dated with high confidence using radiocarbon dating (Shapiro and Hofreiter, 2014). Reconstructing the past population dynamics of bison in this region can help clarify, and improve our understanding of, the contributions of climate change and human presence to megafaunal population decline.

The dataset we use is the same as that in Gill *et al.* (2012) and consists of mtDNA control region sequences from 135 ancient and 17 modern bison samples, with the oldest sample dated 55,182 years before present (BP). We treat sequence sampling dates as known and use the BESP to jointly infer the effective population size trajectory and sampling intensity through time, with *p* = 20 segments, and *p*′ = 12 epochs. Each epoch lasts approximately 5000 years, except for the most recent, which stretches from the present to 450 years BP. We compared our population size estimates to a BSP with 20 population size segments. Both analyses used an HKY+Γ substitution model and a strict molecular clock (see Supplementary Material for further model details).

Fig. 6A shows that sequence sampling has been approximately constant through time, except for the most recent epoch (0–450 years BP), which contains the most samples, and the period 17–22 ka BP, which contains only three samples. This period coincides with the end of the last glacial maximum (LGM), whence fossil material is sparse (Shapiro *et al.*, 2004). The BESP estimates of *N*_*e*_*τ* and *β* through time are shown in blue in Fig. 6B and C, respectively. Estimated effective population size exhibits sustained growth until a population peak around 45 ka BP. This is followed by a population size decline and a population bottleneck around 12 ka BP, with a slight recovery in the recent past.

**Fig. 6:**
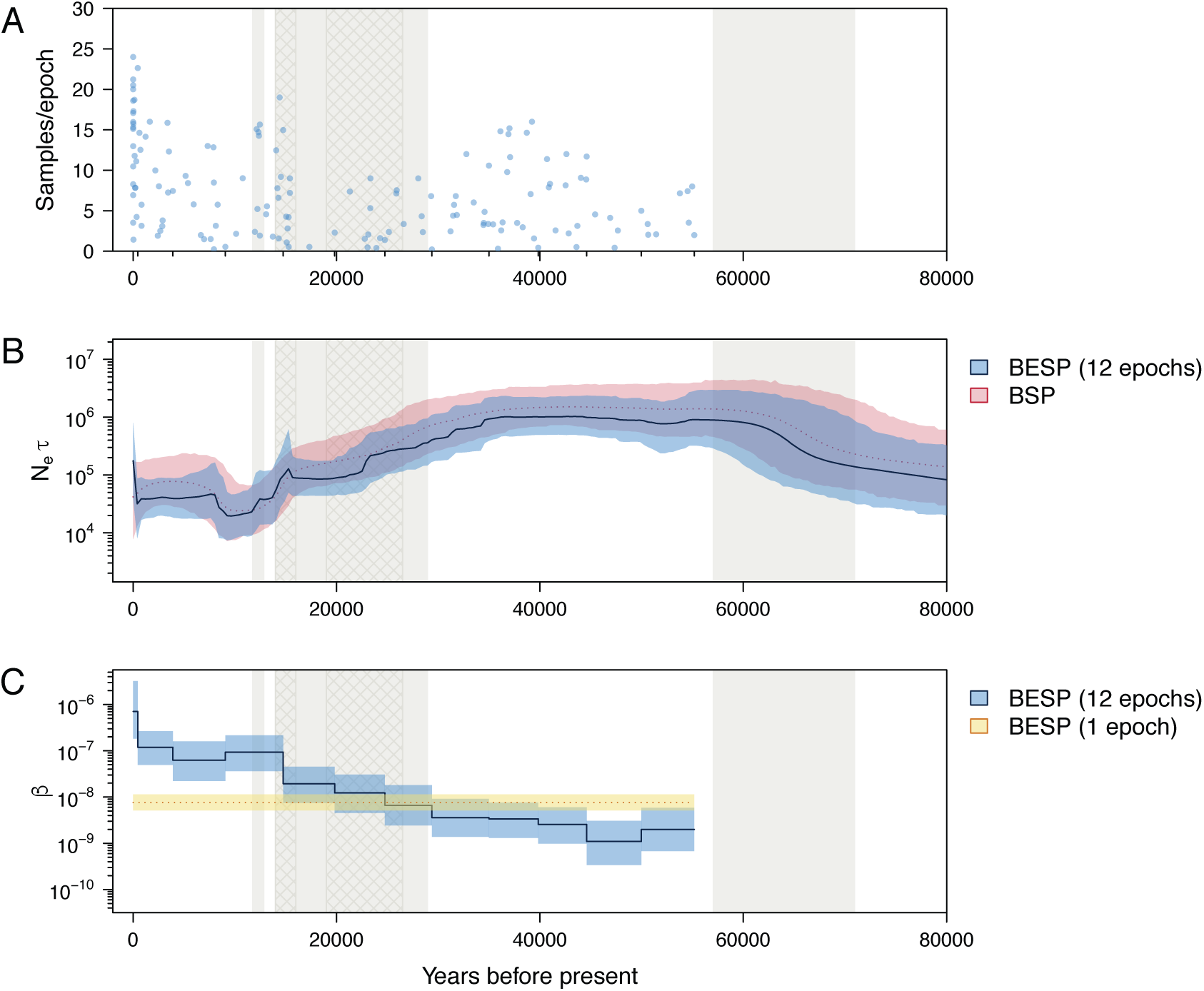
(A) Density of sequence sampling dates through time for the alignment of 152 bison mtDNA sequences that we used. Blue dots indicate stripcharts of individual samples for each sampling epoch. The heights of the stripcharts are equal to the the number of samples in each epoch. Small tick marks on the x-axis represent epoch times. Grey shading indicates cool periods in the Earth’s climate (from the present: Younger Dryas, Marine Isotope Stages (MIS) 2, MIS 4). The two cross hatched areas delimit the time of the last glacial maximum (≈26.5–19 ka BP) and approximate time of substantial human settlement of the Americas (≈16–14 ka BP). (B) Median (solid/dotted line) and 95% highest posterior density (HPD) intervals (shaded areas) for the genetic diversity estimates (*N*_*e*_*τ*) through time. The BESP estimate is shown in blue and the BSP estimate in red. (C) Median (solid line/dotted line) and 95% HPD intervals (shaded areas) of the estimated sampling intensities (*β*) for each sampling epoch. The 12-epoch BESP estimates are in blue and a single-epoch (density-defined) estimate is in yellow.

Both the BESP and BSP infer similar *N*_*e*_*τ* dynamics, with largely overlapping HPD intervals. However, the BESP shows a more rapid and less smooth decline. The BESP recovers a period of stable effective population size around ≈20 ka BP that coincides with the low number of sequences sampled during the LGM. HPD intervals are not notably narrower under the BESP model, likely because phylogenetic uncertainty in this dataset masks any dramatic gains in precision from using the sampling date information.

Estimates of *β*, vary substantially, increasing over four orders of magnitude as time moves from the oldest sample (55 ka BP) towards the present. This contrasts with the limited variation in *β* that was observed in the influenza A virus dataset (Fig. 5C). Thus, this dataset demonstrates how the BESP can be used to detect a strong temporal trend in sampling intensity that requires further exploration.

It is likely that this remarkable increase in sampling intensity is caused by a combination of two factors: (i) sample preservation and successful ancient DNA recovery increases towards the present, and (ii) bison effective population sizes were substantially larger in the past, hence the likelihood of sampling *per-capita* in the past was smaller. There are two notable discontinuous increases in estimated *β*, one at the present (0–450 BP), and one as *N*_*e*_*τ* declines sharply around 15 ka BP. The first is due to the 17 modern sequences in the dataset. The second increase coincides with the period of substantial human settlement of the Americas.

We also investigated a simpler BESP with a single epoch (i.e. density-defined sampling with constant *β* across time). Comparison of the single- and 12-epoch models highlights the significant rise in *β* through time in the latter, and demonstrates that multiple sampling epochs are needed to properly characterise this dataset (Fig. 6C and S9). The single-epoch model generates *N*_*e*_*τ* estimates that are unrealistically high between 15 ka BP and the present, and implies rapid exponential growth in the bison population after the LGM (Fig S9B). This result is an artefact of misspecification of the sampling model: enforcing a constant *β* means that sampling effort in the recent past is greatly underestimated, whilst sampling effort in the distant past is correspondingly overestimated. As a consequence the *N*_*e*_*τ* estimates are biased upwards (downwards) during periods when the sampling intensity is underestimated (overestimated). We conclude that a BESP with a constant *β* is inadequate for this dataset and would promote misleading inferences.

### E. The Information in Sample Timing

We now provide some theoretical basis for why the ESP improves upon the estimates of standard skyline approaches. While sample times are known to provide additional information for demographic inference (Volz and Frost, 2014), their exact contribution has not been quantified. We apply the Fisher information approach from Parag and Pybus (2019) to investigate the benefits of integrating sampling and coalescent events. As in New Approaches, we consider the subtree of 𝒯 that spans the *j*^th^ population size, *N*_*j*_, and contains *s* sampling and *c* coalescent events (see Fig. 1). We use the Fisher information because it delimits the maximum asymptotic precision attainable by any unbiased estimator of *N*_*j*_ (Kay, 1993). This precision defines the inverse of the variance (uncertainty) around that estimator. The Fisher information is computed as the expected second derivative of the log-likelihood (see Materials and Methods).

Popular skyline-based inference methods such as the BSP (Drummond *et al.*, 2005), the skyride (Minin *et al.*, 2008), and the skygrid (Gill *et al.*, 2012), are founded on the coalescent log-likelihood ℒ_*j, c*_, of Eq. (3).

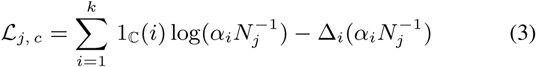

This considers only the *c* coalescent events to be informative about *N*_*j*_. The log-likelihoods specific to each method can be obtained from Eq. (3) by simply altering its population size grouping procedure. The estimates of these approaches are the MLEs of Eq. (3) or some related Bayesian variant. This gives the left side of Eq. (4), which modifies the grouped generalised skyline plot of Strimmer and Pybus (2001) to incorporate the exact times of individual events within that group.

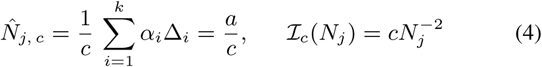

The Fisher information available about *N*_*j*_ from these various skyline-based methods is identical and given by the right side of Eq. (4) (Parag and Pybus, 2019). The maximum precision (minimum variance), around 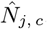, achievable by these approaches is therefore ℐ_*c*_(*N*_*j*_)^*−*1^ (Kay, 1993; Parag and Pybus, 2017).

Next, we define an equivalent log-likelihood for sequence sampling events in Eq. (5). This assumes that only the *s* epochal sampling times are informative and ignores the coalescent events.

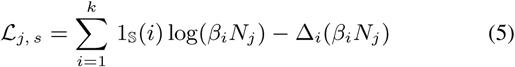

The MLE and Fisher information for this likelihood follow in Eq. (6).

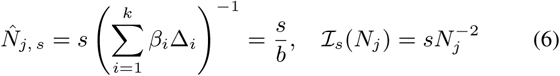

Interestingly, the per event Fisher information 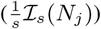 attained by this sampling-event only model is the same as that from any standard skyline method 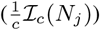. This result formalises and quantifies the assertion in Volz and Frost (2014) that *N*(*t*) can in theory be estimated using only the sampling event times.

Having considered the two information sources separately, we now examine the ESP, which deems both the *s* sampling and *c* coalescent events to be informative. Using Eq. (1) we compute the Fisher information of the *j*^th^ segment, ℐ(*N*_*j*_) (see Materials and Methods). This results in Eq. (7), with 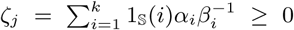 as a grouping factor.

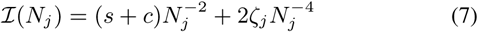

Intriguingly, ℐ(*N*_*j*_) ≥ ℐ_*s*_(*N*_*j*_) + ℐ_*c*_(*N*_*j*_). This means that we gain additional precision by integrating both sampling and coalescent models (the per event information 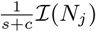 has increased). This extra information comes from the counteracting proportional and inverse dependencies of the two event types. Further, any segment with equal numbers of sampling and coalescent events can now be estimated with at least twice the precision of any standard skyline approach, for the same reconstructed tree 𝒯. Since *n* sampled sequences lead to *n* − 1 coalescent events, and the total Fisher information is 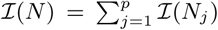, then the overall asymptotic precision across 𝒯 is also roughly, at minimum, doubled.

Eq. (7) explains the improvements in population size inference that the ESP can achieve. However, this improvement may sometimes be clouded by other sources of uncertainty, such as phylogenetic error, and disappears if the sampling times contain no information about population size (in which case the ESP converges to a standard skyline plot). Estimation precision for a given segment depends explicitly on the number of events informing that estimate, i.e. *c* for standard skylines (Eq. (3)), *s* for sampling-events only (Eq. (5)), and *s* +*c* for the ESP (Eq. (1)). This suggests that estimates of *N*_*j*_ should be disregarded when the number of events falling in the *j*^th^ segment is small (if this number is 0 the skyline is unidentifiable as the Fisher information matrix becomes singular (Rothenberg, 1971)). We recommend identifying and excluding such regions from population size estimates as a precaution against overconfident inference.

The log-likelihood of Eq. (1) also provides insight into the statistical power available to infer sampling intensities across time (the *β*_*i*_ parameters). The MLE and Fisher information provided by 𝒯 about *β*_*i*_ over the duration of the *j*^th^ population segment are given in Eq. (8).

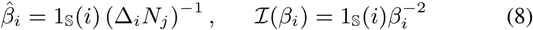

This MLE depends on *N*_*j*_, and thus the two parameters must be jointly estimated (see Materials and Methods for the algorithms that we used to solve this). The Fisher information shows that only intervals ending with sampling events offer the power to estimate a sampling intensity parameter. In our implementation we group the *β*_*i*_ into a smaller number of epochs, so that the power for estimating the sampling intensity during an epoch depends on the total number of sampling events within that epoch. Since, by definition, each epoch contains at least one sampling event, statistical identifiability is guaranteed (Parag and Pybus, 2019). As with population size inferences (discussed above), we recommend ignoring inferences from epochs that contain small numbers of sampling events.

## IV. Discussion

The ESP and its Bayesian implementation (BESP) infer population size history from heterochronous phylogenies and longitudinally-sampled genetic sequences. These methods generalise the skyline approach to include flexible yet tractable models of sequence sampling through time that can more accurately reflect and characterise real-world data collection protocols. This flexible formulation allows the ESP and BESP to serve as tools for the exploration and selection of appropriate time-varying sampling models. This is analogous to how the BSP can be used to select among suitable parametric demographic models for a given dataset.

The improvement in population size inference exhibited by the ESP results from two factors. First, by incorporating sampling time information within an epochal framework we essentially double the number of data points available for inference. As sampling and coalescent events are equally informative (Eq. (4) and Eq. (6)) about population size, we also at least double our best asymptotic estimate precision.

Second, the bias of any coalescent inference method depends on the temporal distribution of its informative events. In standard skyline methods the rate of informative events is inversely dependent on population size, such that periods of large population size possess few coalescent events (resulting in long tree branches), while population bottlenecks feature high event densities. Such skewed distributions can promote inconsistent estimation (Gattepaille *et al.*, 2016). By including sampling events, which cluster in a contrasting way to coalescent events, the ESP achieves more uniform distributions of informative events through time (Fig. 2). This not only reduces bias but also increases temporal resolution, which in turn improves its power to detect and infer rapid population size changes, as seen in both simulated and empirical examples (Fig. 3–Fig. 6).

The ESP was partly inspired by the surveillance and data collection protocols often employed in infectious disease epidemiology. Our assumption that local sampling intensity within an epoch is proportional to population size reflects situations in which sampling is based on availability or convenience, and hence often correlated with the number of infections in an epidemic (Stack *et al.*, 2010). Our inclusion of epochs embodies the expectation that sequence collection rates will likely change discontinuously over time due to fluctuations in funding, resources, and timelines of individual research projects or patient cohorts. Our formulation also allows for external and unpredictable factors that may dramatically alter the sampling effort over an epidemic, such as ‘fog of war’ effects (Viboud *et al.*, 2018).

An analogous situation exists for studies that generate ancient DNA sequences from preserved biological material of different archaeological and geological ages. Specimen preservation and the rate of DNA decay are not only highly dependent on sample age but also on moisture, temperature, and other conditions (Shapiro and Hofreiter, 2014). Thus, while the number of specimens sampled from a given time period might be expected to vary proportionally with species abundance, the constant of proportionality is likely to shift through time. The epoch-based sampling model is sufficiently flexible to capture and extract these types of trends.

Although this flexibility is a benefit of the ESP, we find that biases can result when sampling intensities are defined too rigidly. When an epoch spans a long period of substantial variation in sampling effort, *β* is an estimate of the average sampling intensity over that epoch. If *N* also changes across this epoch then parameter correlations mean that the ESP can overestimate population size in periods where the sampling intensity is underestimated, and vice-versa. This effect is apparent when using the single-epoch model to analyse the Beringian steppe bison dataset (Fig. 6C and S9). This issue possibly underlies the reported biases in previous sampling-aware methods, which all effectively use a single epoch and are based around density-defined models (Karcher *et al.*, 2016).

However, when multi-epoch models are used, the ESP is able to compensate for this bias and expose the vastly different ancient sampling dynamics which underlie this dataset and corroborate previous investigations Shapiro *et al.* (2004). Analysis of the New York influenza epidemic (Fig. 5) highlighted an opposite trend. Here we found that the multi-epoch model offered little advantage over single-epoch formulations, hence providing evidence for a simpler, density-defined description. These results showcase how the ESP can serve as a tool for selecting among various sampling hypotheses and for avoiding model misspecification.

In spite of these benefits, our method has some known limitations. The ESP does not model spatial structure and hence assumes that samples are randomly drawn from a single well-mixed population. Parameter estimates may therefore be biased, if sampling efforts vary across geographic regions. Further, our analysis has relied on having some basic, prior knowledge of how to specify epoch change-points (for example knowing epidemic seasons or understanding practical constraints, as in frequency-defined sampling). If good *a priori* information is unavailable and epoch times are set arbitrarily, biases can result as we may have periods over which the ESP is too rigidly formulated (akin to the single-epoch bias). In these cases we recommend distributing sampling events evenly among epochs to guard against this type of misspecification.

The ESP differs from previous approaches that use parametric sampling models (Karcher *et al.*, 2016; Volz and Frost, 2014). This mirrors the distinction between skyline plot methods and coalescent estimators of parametric demographic functions (Parag and Pybus, 2017). Karcher *et al.* (2016), for example, used a non-linear sampling rate model of form *e*^*γ*0^ *N*(*t*)^*γ*1^, with *γ*_0_ and *γ*_1_ as parameters to be inferred. While such formulations do not model the same range of sampling behaviours as the ESP, they can provide specific biological insights (e.g. *γ*_1_ informs about sample clustering) if the true (unknown) sampling rate lies within their functional class. The ESP, by providing insight into what types of parametric hypotheses might be supported by a given dataset, can complement these approaches.

The sampling intensity, *β*, inferred by the ESP can be used to reconstruct the absolute sampling rate, *βN*. Practically, *β* measures how quickly new sequences accumulate relative to the effective population size (i.e. ‘per capita’). It has units of [time^*−*2^]. Since *N* has dimensions of [time] (measured in the units of the time-scaled genealogy) then the ESP directly infers changes in the rate of collecting samples per genealogical time unit. The separation of *β* and *N* is important, as it disaggregates the relative contributions of each time-varying unknown. Further, as *β* modulates a Poisson process, then over an infinitesimal period it defines a piecewise-constant sampling probability that is analogous to (but not equal to) the sampling model used in phylogenetic birth-death skyline methods (Stadler *et al.*, 2013).

As sequence data becomes more prevalent, heterochronous sampling design will play an increasingly important role in phylodynamics (Ho and Shapiro, 2011; Parag and Pybus, 2019). Continuing improvements in infectious disease monitoring and sequencing will result in richer and more diverse epidemiological data (Baele *et al.*, 2017), while ongoing advances in techniques for isolating and generating ancient DNA will lead to strengthened molecular evolution datasets. We hope that the ESP will prove useful in exploiting and exploring such data and help inform the debates surrounding sequence sampling protocol design and misspecification.

## V. Materials and Methods

### A. Deriving the Epoch Sampling Skyline Plot

Here we construct the log-likelihood for the ESP (Eq. (1)), and derive its population size MLE (Eq. (2)) and Fisher information (Eq. (7)). Let the *j*^th^ piecewise-constant segment of a sampled-coalescent process have unknown population size *N*_*j*_, and duration 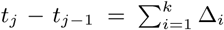. We assume that this segment consists of *k* ≥ 1 event intervals, the *i*^th^ of which has duration Δ_*i*_. If this interval ends in a sampling (coalescent) event, then 1_𝕊_(*i*) = 1(0), and 1_ℂ_(*i*) = 0(1). The coalescent lineage factors, and sampling intensities, for the *i*^th^ interval are respectively *α*_*i*_ and *β*_*i*_. Fig. 1 clarifies this notation for a simple reconstructed coalescent genealogy (tree), 𝒯, over this segment. Standard skyline approaches model coalescent events as the outputs of a Poisson process with rate 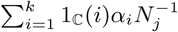, but ignore sampling events. The ESP assumes that sampling events are also produced by a Poisson process, with rate 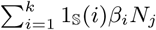. The result is a piecewise-constant dual-type Poisson process, with combined event rate *λ*(*t*) as in Eq. (9).

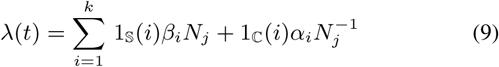

We construct the combined Poisson log-likelihood function for the *j*^th^ segment, ℒ_*j*_: = log P(𝒯 | *N*_*j*_, {*β*_*i*_}), as in Eq. (10) (Snyder and Miller, 1991; Parag and Pybus, 2018).

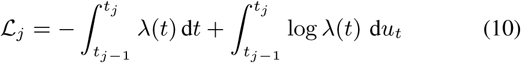

The total log-likelihood over all *p* segments of 𝒯 is 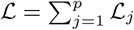. For now we only focus on the set of *N*_*j*_ unknowns in this log-likelihood (we discuss the power to estimate {*β*_*i*_} in the next section). In Eq. (10), d*u*_*t*_ = 1 at event times, and 0 otherwise, so that the second integral is a sum over interval end-points. Eq. (1) is derived by splitting the integrals in Eq. (10) over the *k* intervals. Note that defines population size change-points at (irregular) event times. This contrasts with the approach of Karcher *et al.* (2016), where change-point times are regular, predefined and do not depend on the temporal event distribution. One advantage of our formulation is that we always have at least one event informing on each *N*_*j*_ parameter. This results in a non-singular Fisher information matrix, which guarantees the statistical identifiability of the ESP (Rothenberg, 1971) (Parag and Pybus, 2019).

The skyline estimator that we propose is the grouped MLE of Eq. (1). This solves 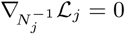 when *s* ≥ *c*, and leads to the quadratic expression in *N*_*j*_ given in Eq. (11).

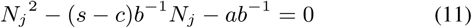

Here ∇_*x*_ is the first partial derivative with respect to *x*, while 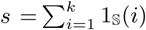, and 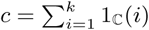 count the total number of sampling and coalescent events falling in the *j*^th^ segment of 𝒯. If *s* < *c* then 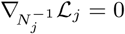 must be computed, and then inverted. This gives Eq. (12), which is a quadratic in 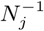.

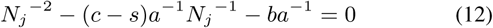

This conditional MLE approach is needed to avoid singularities in cases when either *s* = 0, or *c* = 0, and to keep population sizes positive. The roots of these quadratics form Eq. (2).

The Fisher information of the ESP with respect to *N*_*j*_, is defined 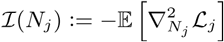, with 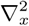 as the second partial derivative (Kay, 1993). The expectation is taken across the event intervals, Δ_*i*_. Applying this to Eq. (1), we obtain Eq. (13).

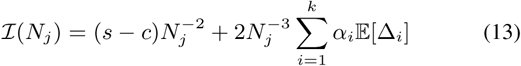

Note that we can replace ℒ_*j*_ with ℒ_*c, j*_ or ℒ_*s, j*_ in the above definition, to also recover Eq. (4) and Eq. (6), the Fisher information stemming from only the coalescent and sampling events, respectively. The expectation in Eq. (13) conditions on the type of event in each interval i.e. 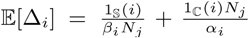. Expanding 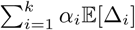 we get 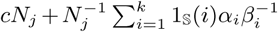. Substituting this into Eq. (13) simplifies to Eq. (7), which when *s* ≈ *c* reveals a minimum ℐ(*N*_*j*_) of 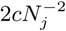. This is twice the value obtained in Eq. (4), and shows the marked improvement in estimate precision that results from including sampling events.

Lastly, we comment on how ESP population size estimates relate to those in Eq. (4) and Eq. (6). We group our skyline over the entire tree so that there is only a single population size to estimate, *N*_1_. This is equivalent to a Kingman coalescent assumption (i.e. constant population size). Since the number of coalescent and sampling events are always roughly the same then we can use the *s* ≈ *c* solution of Eq. (2), and the MLEs from Eq. (4) and Eq. (6) to derive 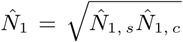. If we think of the true population size, *N*(*t*), as being continuously time-varying, then standard skylines estimate its harmonic mean with 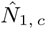 (Pybus *et al.*, 2000). Similarly, 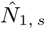 estimates the arithmetic mean of *N*(*t*). The ESP is then the geometric mean of these two mean estimators, and hence smooths the individual population size estimates from Eq. (4) and Eq. (6).

### B. Estimating the Epoch Sampling Intensities

We now define our epochal sampling model, characterise the power of the ESP for estimating sampling intensities, and present algorithms to compute the ML estimates of these sampling intensities. We assume a total of *p*′ epochs, spanning the duration of the first (most recent) to last (most ancient) sampling event (time increases into the past). This is the period over which non-zero sampling effort is assumed. Within each epoch, the sampling intensities of each interval are the same, and epoch times coincide with sample times. This results in a piecewise-constant, time delimited, longitudinal sampling intensity. We first consider the most flexible, naïve epoch model, in which each interval is treated as a new epoch. For the *j*th segment, this means there are *k* sampling unknowns, {*β*_*i*_}. The MLE, 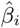, is the solution to 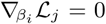. The Fisher information that 𝒯 contains about 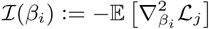.

Applying these to Eq. (1) gives Eq. (8), the MLE and Fisher information of *β*_*i*_ during the *j*^th^ population segment. Two key observations emerge: (i) 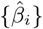 depends on 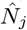, and (ii) we only have power to estimate sampling intensities in intervals that contain sampling events (ℐ(*β*_*i*_) = 0 | *i* ∈ ℂ). Point (ii) suggests that if *i*′ ∈ 𝕊 and *i*′ + 1 ∈ ℂ then we should assume either *β*_*i*′+1_ = 0 or *β*_*i*′+1_ = *β*_*i*′_, to ensure identifiability. We can resolve (ii) by grouping our sampling intensities (similar to how we group over *N*_*j*_) so that there are only *p*′ distinct epochs. Within these epochs there is only one sampling intensity parameter, and there is always at least one sampling event, guaranteeing identifiability (the Fisher information with respect to grouped *β*_*i*_ is non-singular (Rothenberg, 1971)). The minimum variance around these per-epoch estimates of sampling intensity is then related to the sum of the ℐ(*β*_*i*_) comprising the epoch. For example, if there is 1 epoch over the *j*^th^ segment, with unknown intensity *β*_*j*_, then 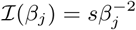.

Thus the ESP contains power to estimate (sensibly) flexible sampling intensity changes through time. Computing these estimates, and hence resolving (i), requires joint inference of the population size and sampling intensity parameters. For ML inference, we achieve this with a simple iterative algorithm. Let *β* and *N* be the *p*′ and *p* element vectors of unknowns that we want to estimate. We draw an initial 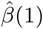 from a wide uniform distribution and then compute the conditional estimate 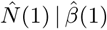 using Eq. (2). We substitute this into Eq. (8) to get 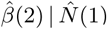. Repeating this procedure iteratively yields the desired joint MLEs, 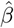 and 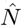, usually within 100 steps (it does not require tuning and is robust to the initial 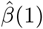). This algorithm, and the above ML solutions, are all implemented in Matlab and available at https://github.com/kpzoo/epoch-sampling-skyline-plot.

### C. The Bayesian Epoch Sampling Skyline Plot

Here we extend the BSP (Drummond *et al.*, 2005) to incorporate the epochal sampling model defined in the previous section. Given a genealogy 𝒯, a set of *p* segment sizes, *K* = {*k*_1_, *k*_2_, …, *k*_*p*_}, counting the numbers of events (coalescent/sampling) in each piece-wise population size segment, and a set of *p*′ epoch sizes, 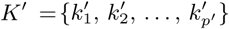, counting the sampling events in each epoch, we can compute the likelihood *f* (𝒯|*N, K, β, K*′) from Eq. (1). Applying Bayes’ theorem yields the joint posterior distribution of *N, β* and *K* given in Eq. (14).

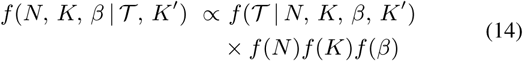

We obtain the Bayesian ESP (i.e. BESP) by sampling from this posterior using standard MCMC proposal distributions. Eq. (14) features priors on the population size vector, *N*, its grouping parameter (the number of events in each population size segment), *K*, and the sampling intensity vector, *β*. We have assumed that *p, p*′ and the epoch grouping parameter, *K*′, are all specified *a priori*, which reflects the belief that we generally have a reasonable idea of the timescale over which sampling intensities vary. This assumption could in theory be relaxed by sampling epoch sizes (*K*′) within BEAST2.

We impose the same smoothing prior on *N* as in the BSP. This assumes that neighbouring effective population size segments are autocorrelated, and implements this by drawing *N*_*j*_ from an exponential distribution with a mean equal to *N*_*j*−1_ (i.e. 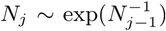 for 2 ≤ *j* ≤ *p*) and a Jeffreys prior on *N*_1_ (Drummond *et al.*, 2005). Since we expect sampling efforts to change discontinuously we do not assume that neighbouring sampling intensities are autocorrelated, and place independent and identical priors on each *β*_*i*_. It is trivial to relax this assumption and apply different priors to each *β*_*i*_, e.g. if *a priori* information is available about changes in sampling effort through time. This is analogous to the recent approach in Karcher *et al.* (2019), which embeds time-varying external covariates within the sampling process.

Our BESP implementation also contains some practical adjustments. We constrain the minimum segment duration for both population size segments and sampling epochs to be above some threshold *E*. This guards against zero-length segments or epochs, which can result if too many sampling events coincide in time or if phylogenies contain bursts of branching events. Further, we constrain segments and epochs to contain at least two informative events each, to safely ensure identifiability. The BESP is implemented as a BEAST2.6 (Bouckaert *et al.*, 2019) package and uses *f* (𝒯 |*N, K, β, K*′) as a tree-prior for Bayesian phylogenetic analysis. This allows the BESP, in conjunction with existing substitution and clock models, to jointly infer changes in effective population size and sampling intensity directly from sequence data, while incorporating phylogenetic uncertainty. The BESP package is available at https://github.com/laduplessis/besp and raw data, workflows, XML files and additional figures for the simulations and empirical analyses presented above are available at https://github.com/laduplessis/BESP_paper-analyses.

## VI. Acknowledgments

We thank Edward Holmes, Beth Shapiro, Cecile Viboud and Amanda Perofsky for access to and advice on the empirical datasets. This work was supported by the European Research Council under the European Commission Seventh Framework Programme (FP7/2007-2013)/European Research Council grant agreement 614725-PATHPHYLODYN. We also acknowledge support from the Oxford Martin School and the UK MRC and Department for International Development under grant reference MR/R015600/1.

## Appendix

### A. Bayesian Implementation Simulation Study

#### 1) Simulations and Inferences

We used the `phylodyn` **R** package (Karcher *et al.*, 2017) to simulate heterochronously sampled coalescent trees under five population size trajectories, *N*(*t*), given in table S1. Trees were simulated with approximately 500 samples, sampled between *t* = 0 and *t*: = min{48,⌊*t*_10_⌋} where *t*_10_ ={*t*: *N*(*t*) ≤ 10}. The sampling period was divided into 24 equally-spaced epochs, with the sampling intensity (*β*) during each epoch set such that an approximately equal number of samples is drawn from each epoch. We used this procedure to simulate 100 replicate trees for each population size trajectory.

Inferences were performed for each simulated tree using the BESP implementation in BEAST2 v2.5 (Bouckaert *et al.*, 2019). To estimate *N* we set *p* = 100, and grouped equal numbers of events (coalescent/sampling) so that each population size segment had roughly 10 informative events i.e. *k*_*j*_ ≈ 10 for all *j*. To estimate *β* we set *p*′ = 24, with the boundaries of sampling epochs chosen to coincide with the time of the first sampling event after each sampling epoch change (going into the past). Because each simulation contains ≈500 samples, in practice this resulted in sampling intensity change-point times that were very close to the true epoch times used in the simulations. In all analyses *N* and *β* were jointly estimated, and all other parameters (including the tree) were fixed to the truth.

MCMC chains were run for 20 million iterations and parameters sampled every 20,000 iterations. Convergence was checked by calculating the effective sample size (ESS) of parameters using the `coda` R package (Plummer *et al.*, 2006) after discarding 10% of samples as burn-in. All parameters had ESS values greater than 200 in all but 3 analyses, all computed on trees simulated under a boom-bust population size trajectory. The MCMC traces of these 3 analyses were checked manually using the program Tracer v1.7 (Rambaut *et al.*, 2018) to confirm convergence. Workflows for the Bayesian simulation study are availabe at https://github.com/laduplessis/BESP_paper-analyses.

#### 2) Summary Statistics

Let *θ*_*i*_ be the true values of a vector of parameters, where *θ*_*i*_ is defined over the period [*t*_*i*−1_, *t*_*i*_), with *t*_*i*_ > *t*_*i*−1_ for 1 ≤ *i* ≤ *p*. If 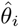 is the median *a posteriori* estimate of *θ*_*i*_ and 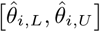 is the HPD interval we calculate the following statistics:

i. *Mean relative bias:*

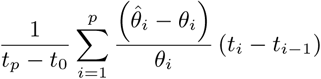
ii. *Mean relative HPD interval width:*

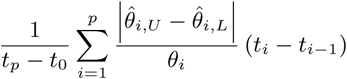
iii. *Mean coverage:*

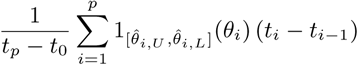

We use the above three statistics to calculate summary statistics for *N* (across population size segments) and *β* (across sampling epochs), between the most recent and most ancient samples. Since the true population size trajectory, *N*(*t*) varies across each segment we use the harmonic mean of *N*(*t*) within a segment as the true parameter value. This is a lower bound on the true parameter estimated by the BESP, with the true estimated parameter being a product of the harmonic and arithmetic means (see equation 2 in the main text). By the simulation design *β* does not vary substantially over each sampling epoch and we simply use the true *β* to compute its summary statistics.

### B. Case Study 1: Seasonal Human Influenza

We analysed a subset of the 687 human influenza A/H3N2 HA gene sequences sampled in New York state between 1993 and 2005 that were used in the analyses presented in Rambaut *et al.* (2008). We removed all sequences sampled during the 1992/1993 influenza season, as only isolates from the second half of this season were included in the dataset. The remaining dataset contains 637 sequences representing 12 complete influenza seasons (1993/1994–2004/2005). The HA alignment is the same as that in Rambaut *et al.* (2008) but with 50 sequences removed, comprises the coding regions of the HA segment and spans 1,698 bp.

We used an SRD06 substitution model (Shapiro *et al.*, 2006), that allows for different rates of evolution of the first and second codon positions relative to the third, and uses an HKY nucleotide substitution model with Γ-distributed rate heterogeneity for both partitions. We further utilised an uncorrelated lognormal relaxed clock model (Drummond *et al.*, 2006) to allow for variations in the overall rate of molecular evolution across branches in the tree. Default priors were used for the nucleotide substitution model and an informative lognormal prior was placed on the mean molecular clock rate, with a mean of 6 × 10^*−*^3 s/s/y and *σ* = 0.1. As a tree-prior we used either the BESP or BSP, with three different configurations:

i. **BESP (12 epochs)**: *p* = 40, *p*′ = 12, *k*_*j*_ ≥ 2, *t*_*j*_ − *t*_*j*−1_ > 0.08 years (≈ 1 month) for population size segments and sampling epochs. Sampling epoch group sizes (*K*′) and change-point times are fixed as in table S2.
ii. **BESP (1 epoch)**: *p* = 40, *p*′ = 1, *k*_*j*_ ≥ 2, *t*_*j*_ − *t*_*j*−1_ > 0.08 years (≈ 1 month) for population size segments. *β* is constant between 2005.25 and 1993.87 and 0 between 1993.87 and the time of the most recent common ancestor.
iii. **BSP**: *p* = 40, *k*_*j*_ ≥2, *t*_*j*_ − *t*_*j*−1_ > 0.08 years (≈1 month) for population size segments.

**TABLE S1:** Population size trajectories used in the Bayesian implementation simulation study.

|  |  |  |
| --- | --- | --- |
| Constant size | $N(t) = 100$ | |
| Bottleneck | $N(t) = \begin{cases} 100 \\ 10 \end{cases}$ | $t \leq 10, t \geq 20$<br>$10 < t < 20$ |
| Boom-bust | $N(t) = \begin{cases} 500 \exp(t - 2) \\ 500 \exp(2 - t) \end{cases}$ | $t \leq 2$<br>$t > 2$ |
| Cyclical boom-bust | $N(t) = \begin{cases} 200 \exp(-5(t \bmod 10)) \\ 200 \exp(5[(t \bmod 10) - 10]) \end{cases}$ | $t \bmod 10 \leq 5$<br>$t \bmod 10 > 5$ |
| Logistic growth and decline | $N(t) = \begin{cases} 10 + \frac{190}{1 + \exp(2[3 - (t \bmod 12)])} \\ 10 + \frac{190}{1 + \exp(2[3 + (t \bmod 12) - 12])} \end{cases}$ | $t \bmod 12 \leq 6$<br>$t \bmod 12 > 6$ |

To aid convergence we computed an initial maximum-likelihood genetic distance tree using RAxML v8 (Stamatakis, 2014) under a general time reversible nucleotide substitution model with G-distributed rate heterogeneity. We then used TreeTime (Sagulenko *et al.*, 2018) to construct a time-calibrated initial tree with branch lengths in years. The models we use differ from the model used in Rambaut *et al.* (2008) only in the prior distributions, initial tree and bounds on the number of events in each segment/epoch and segment/epoch lengths. Rambaut *et al.* (2008) only used the BSP (with no minimum group size), set no explicit priors for clock and substitution model parameters and imposed no bounds on segment parameters.

All analyses were performed in BEAST v2.5 (Bouckaert *et al.*, 2019). For each model, we computed 7 independent MCMC chains of 200 million iterations and sampled parameters and trees every 10,000 iterations. We used custom R scripts to combine chains after discarding 30% of samples as burn-in. The combined chain was thinned by a factor of 3 and convergence was checked by calculating the ESS of parameters using the `coda` R package (Plummer *et al.*, 2006). The marginal posterior estimates of *N*(*t*) were obtained by discretising the *N*_*j*_ parameters over an even grid of 80 cells, between 2005.25 and 1992.25 using a custom R script. To compute the maximum clade credibility (MCC) tree, we used the program `logcombiner` to combine the 7 sets of tree samples after discarding 30% of samples and thinning by a factor of 3. The program `treeannotator` was then used to compute the MCC tree of the resulting posterior tree distribution. Workflows for the seasonal human influenza case study are availabe at https://github.com/laduplessis/BESP_paper-analyses.

### C. Case Study 2: Steppe Bison

We analysed 152 bison mtDNA control region sequences of 602 bp, dating from the present to 55,182 years BP. The alignment is the same as the one used in Gill *et al.* (2012). In the analyses we used an HKY substitution model (Hasegawa *et al.*, 1985) and a strict clock model. Default priors were used for the nucleotide substitution model and a uniform prior on (0, ∞) was placed on the molecular clock rate. All analyses were started from a random constant population size coalescent tree. As a tree-prior we used either the BESP or BSP, with three different configurations:

i. **BESP (12 epochs)**: *p* = 20, *p*′ = 12, *k*_*j*_ ≥2, *t*_*j*_ −*t*_*j*−1_ > 100 years for population size segments and sampling epochs. Sampling epoch group sizes (*K*′) and change-point times are fixed as in table S3.
ii. **BESP (1 epoch)**: *p* = 20, *p*′ = 1, *k*_*j*_≥ 2, *t*_*j*_ −*t*_*j*−1_ > 100 years for population size segments. *β* is constant between the present and 55,182 years BP and 0 between 55,182 years BP and the time of the most recent common ancestor.
iii. **BSP**: *p* = 20, *k*_*j*_ ≥2, *t*_*j*_ − *t*_*j*−1_ > 100 years for population size segments.

**TABLE S2:** Sampling epochs used for the 12-epoch BESP on the influenza A/H3N2 dataset.

| Season | Samples | Start | End | Years |
| --- | --- | --- | --- | --- |
| 2004/2005 | 80 | 2005.25 | 2004.73 | 0.52 |
| 2003/2004 | 85 | 2004.73 | 2003.64 | 1.09 |
| 2002/2003 | 16 | 2003.64 | 2003.07 | 0.57 |
| 2001/2002 | 84 | 2003.07 | 2001.91 | 1.16 |
| 2000/2001 | 1 | 2001.91 | 2001.08 | 0.83 |
| 1999/2000 | 70 | 2001.08 | 1999.76 | 1.32 |
| 1998/1999 | 78 | 1999.76 | 1998.85 | 0.91 |
| 1997/1998 | 49 | 1998.85 | 1997.92 | 0.93 |
| 1996/1997 | 51 | 1997.92 | 1996.88 | 1.04 |
| 1995/1996 | 25 | 1996.88 | 1995.92 | 0.96 |
| 1994/1995 | 49 | 1995.92 | 1994.59 | 1.33 |
| 1993/1994 | 49 | 1994.59 | 1993.87 | 0.72 |

**TABLE S3:** Specification of the sampling epochs for the bison dataset.

| Epoch | Samples | Start (BP) | End (BP) | Years |
| --- | --- | --- | --- | --- |
| 1 | 24 | 0 | 450 | 450 |
| 2 | 16 | 450 | 3,903 | 3,453 |
| 3 | 13 | 3,903 | 9,068 | 5,165 |
| 4 | 19 | 9,068 | 14,753 | 5,685 |
| 5 | 9 | 14,753 | 19,815 | 5,062 |
| 6 | 9 | 19,815 | 24,752 | 4,937 |
| 7 | 9 | 24,752 | 29,377 | 4,625 |
| 8 | 12 | 29,377 | 34,987 | 5,610 |
| 9 | 16 | 34,987 | 39,836 | 4,849 |
| 10 | 12 | 39,836 | 44,598 | 4,762 |
| 11 | 5 | 44,598 | 49,975 | 5,377 |
| 12 | 8 | 49,975 | 55,182 | 5,207 |

All analyses were performed in BEAST v2.5 (Bouckaert *et al.*, 2019). For each model, we computed 3 independent MCMC chains of 200 million iterations and sampled parameters and trees every 10,000 iterations. We used a custom R script to combine chains after discarding 10% of samples as burn-in. The combined chain was thinned by a factor of 3 and convergence was checked by calculating the ESS of parameters using the `coda` R package (Plummer *et al.*, 2006). The marginal posterior estimates of *N*(*t*) were obtained by discretising the *N*_*j*_ parameters over an even grid of 200 cells, between the present and 80 ka BP using a custom R script. Workflows for the steppe bison case study are availabe at https://github.com/laduplessis/BESP_paper-analyses.

**Fig. S1:**
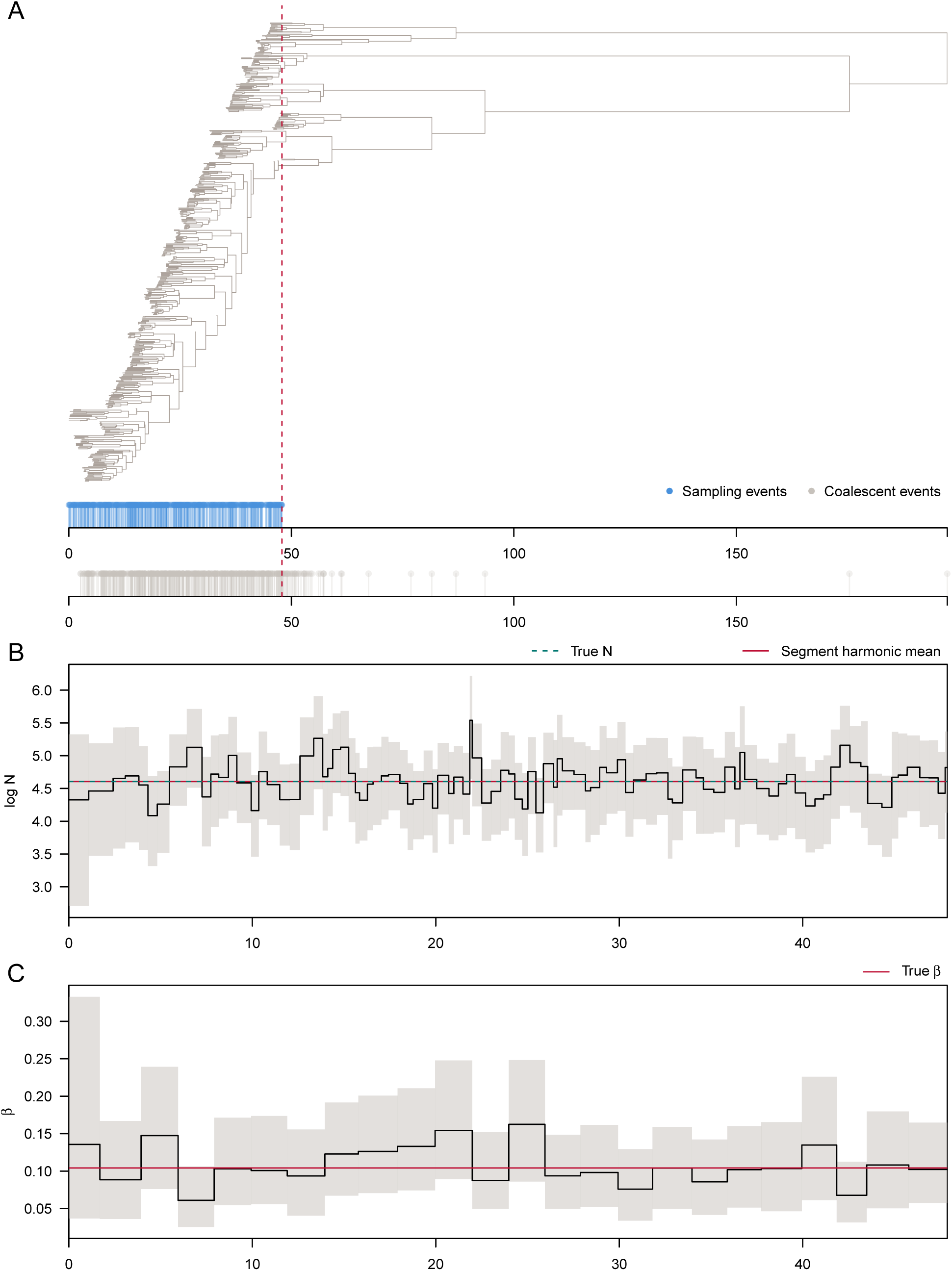
(A) Example of one of the 100 replicate trees simulated under the constant size demographic scenario. Sampling (blue) and coalescent (grey) events are shown below. The red dashed line indicates the time of the most ancient sample. (B) Median (solid black line) and HPD intervals (shaded areas) for the effective population size (*N*) estimates between the most recent and most ancient samples. The dashed green line shows the true *N*-trajectory used to simulate the tree in A and the red line the harmonic mean of the true *N* during each segment. (C) Median (solid black line) and HPD intervals (shaded areas) for the sampling intensity (*β*) estimates for each sampling epoch. The red line shows the true *β* used to simulate the tree in A.

**Fig. S2:**
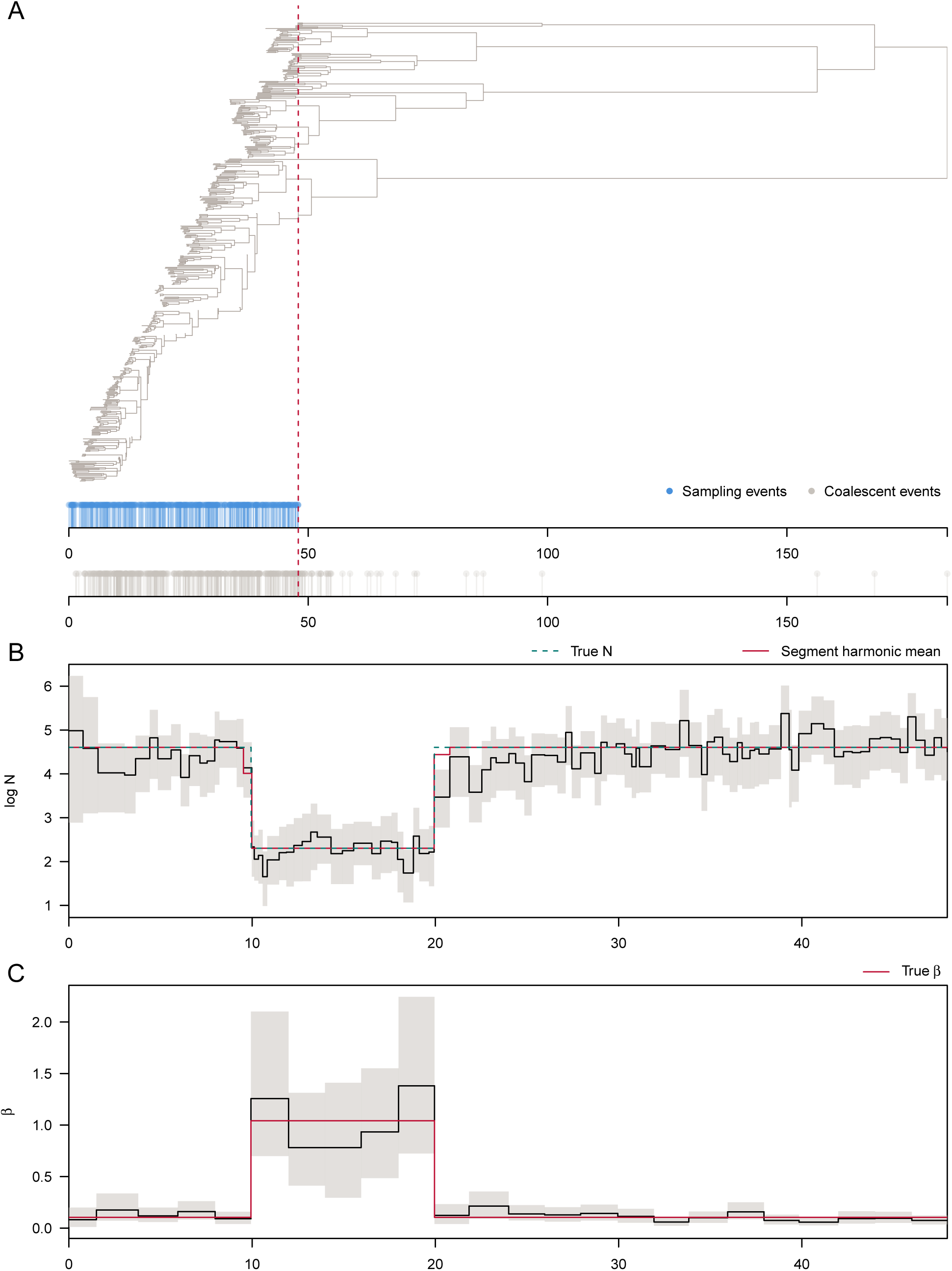
(A) Example of one of the 100 replicate trees simulated under the bottleneck demographic scenario. Sampling (blue) and coalescent (grey) events are shown below. The red dashed line indicates the time of the most ancient sample. (B) Median (solid black line) and HPD intervals (shaded areas) for the effective population size (*N*) estimates between the most recent and most ancient samples. The dashed green line shows the true *N*-trajectory used to simulate the tree in A and the red line the harmonic mean of the true *N* during each segment. (C) Median (solid black line) and HPD intervals (shaded areas) for the sampling intensity (*β*) estimates for each sampling epoch. The red line shows the true *β* used to simulate the tree in A.

**Fig. S3:**
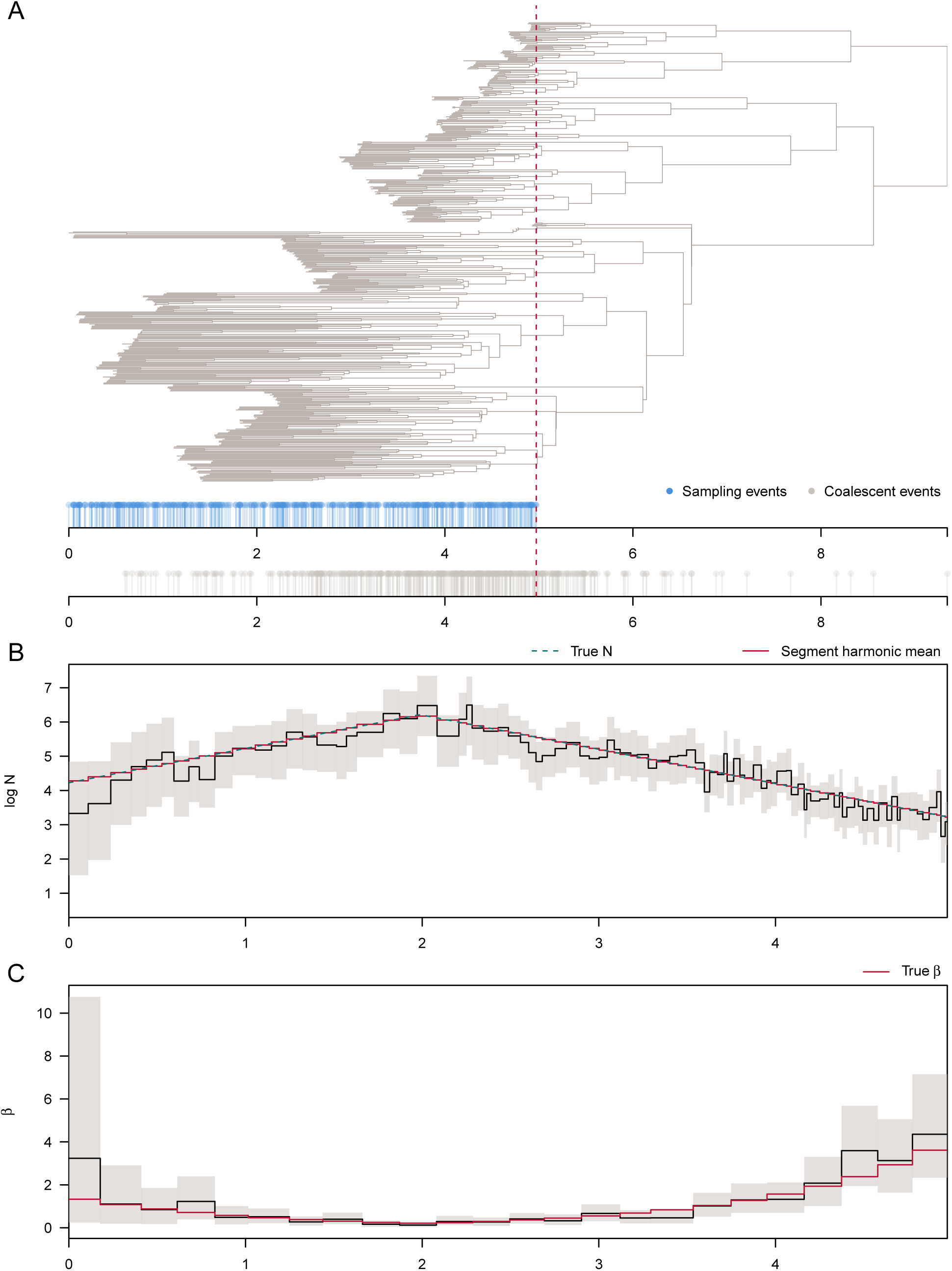
(A) Example of one of the 100 replicate trees simulated under the boom-bust demographic scenario. Sampling (blue) and coalescent (grey) events are shown below. The red dashed line indicates the time of the most ancient sample. (B) Median (solid black line) and HPD intervals (shaded areas) for the effective population size (*N*) estimates between the most recent and most ancient samples. The dashed green line shows the true *N*-trajectory used to simulate the tree in A and the red line the harmonic mean of the true *N* during each segment. (C) Median (solid black line) and HPD intervals (shaded areas) for the sampling intensity (*β*) estimates for each sampling epoch. The red line shows the true *β* used to simulate the tree in A.

**Fig. S4:**
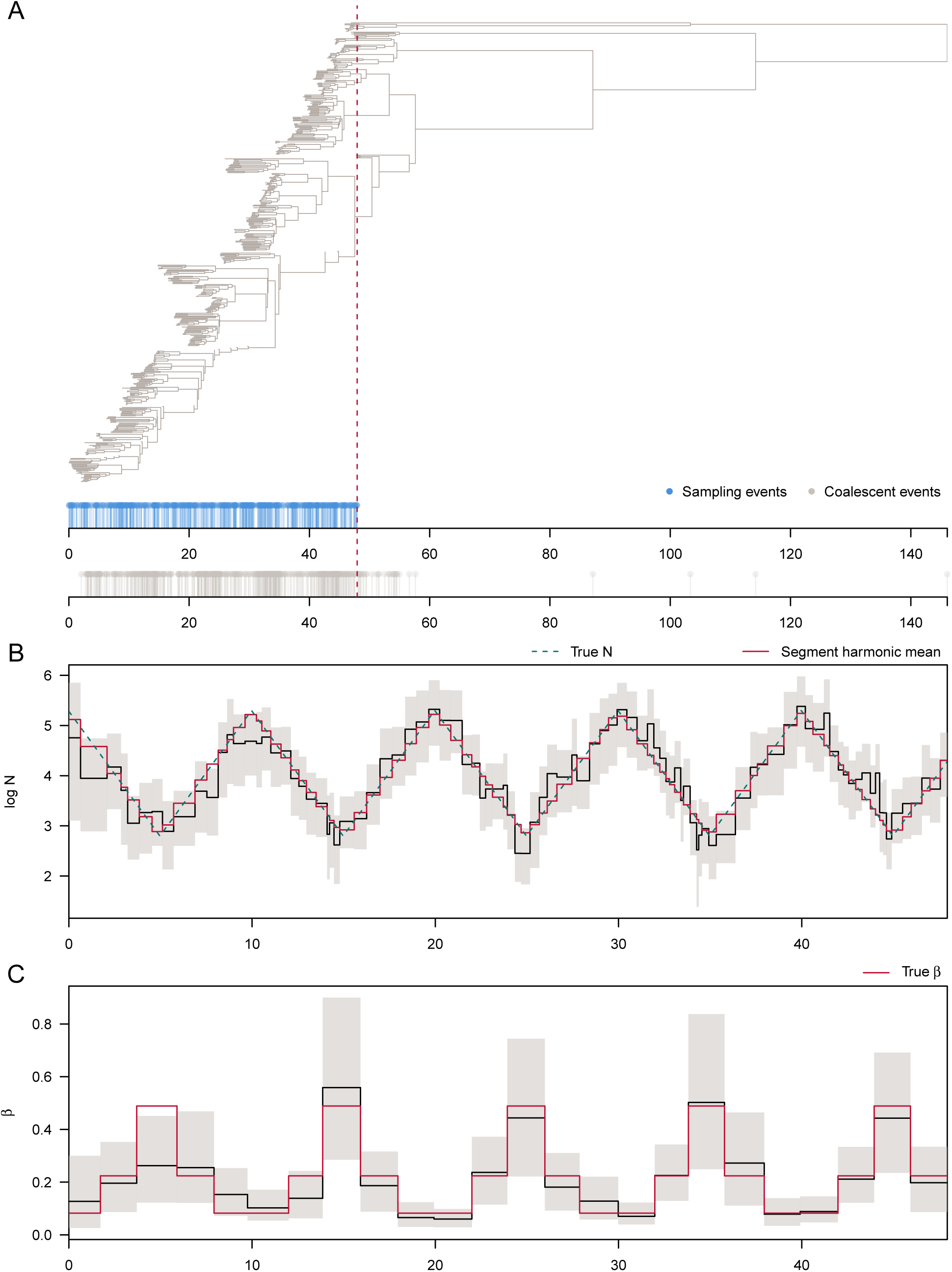
(A) Example of one of the 100 replicate trees simulated under the cyclical boom-bust demographic scenario. Sampling (blue) and coalescent (grey) events are shown below. The red dashed line indicates the time of the most ancient sample. (B) Median (solid black line) and HPD intervals (shaded areas) for the effective population size (*N*) estimates between the most recent and most ancient samples. The dashed green line shows the true *N*-trajectory used to simulate the tree in A and the red line the harmonic mean of the true *N* during each segment. (C) Median (solid black line) and HPD intervals (shaded areas) for the sampling intensity (*β*) estimates for each sampling epoch. The red line shows the true *β* used to simulate the tree in A.

**Fig. S5:**
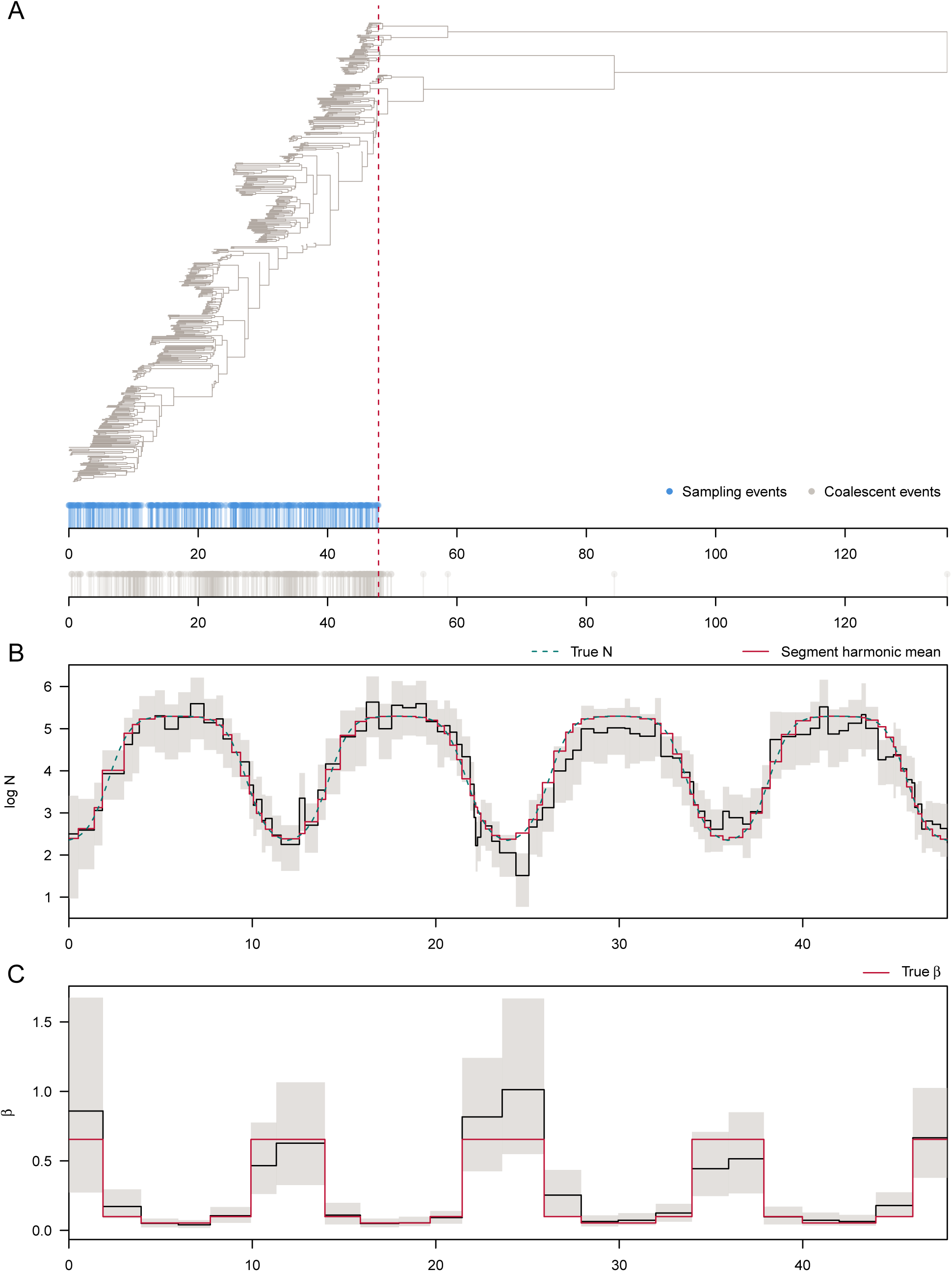
(A) Example of one of the 100 replicate trees simulated under the logistic growth and decline demographic scenario. Sampling (blue) and coalescent (grey) events are shown below. The red dashed line indicates the time of the most ancient sample. (B) Median (solid black line) and HPD intervals (shaded areas) for the effective population size (*N*) estimates between the most recent and most ancient samples. The dashed green line shows the true *N*-trajectory used to simulate the tree in A and the red line the harmonic mean of the true *N* during each segment. (C) Median (solid black line) and HPD intervals (shaded areas) for the sampling intensity (*β*) estimates for each sampling epoch. The red line shows the true *β* used to simulate the tree in A.

**Fig. S6:**
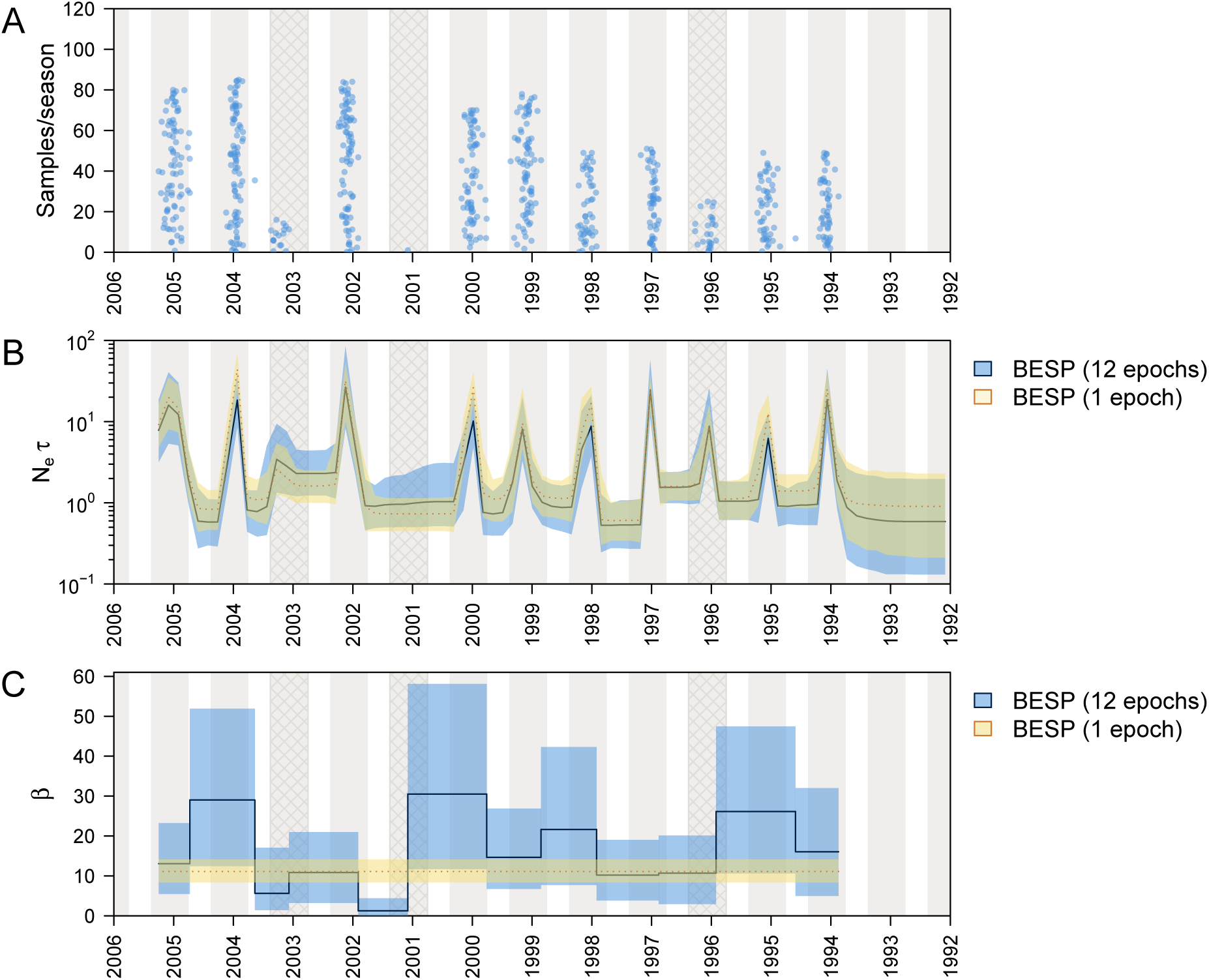
(A) Density of sequence sampling dates through time for the alignment of 637 A/H3N2 HA sequences from NY state that we analysed. Blue dots indicate stripcharts of individual samples for each season. The stripchart heights give the number of samples in each season. Grey shading indicates the approximate period of influenza observation in New York state during each season (epidemiological week 40, to week 20 in the next year). Cross-hatched seasons are those where A/H3N2 was not the dominant influenza virus subtype. (B) Median (solid/dotted line) and 95% highest posterior density (HPD) intervals (shaded areas) for the genetic diversity estimates (*N*_*e*_*τ*) through time. The 12-epoch BESP estimate is shown in blue and the single-epoch BESP estimate is in yellow. (C) Median (solid line/dotted line) and 95% HPD intervals (shaded areas) of the estimated sampling intensities (*β*) for each sampling epoch. The 12-epoch BESP estimates are shown in blue and a single-epoch (density-defined) estimate is in yellow.

**Fig. S7:**
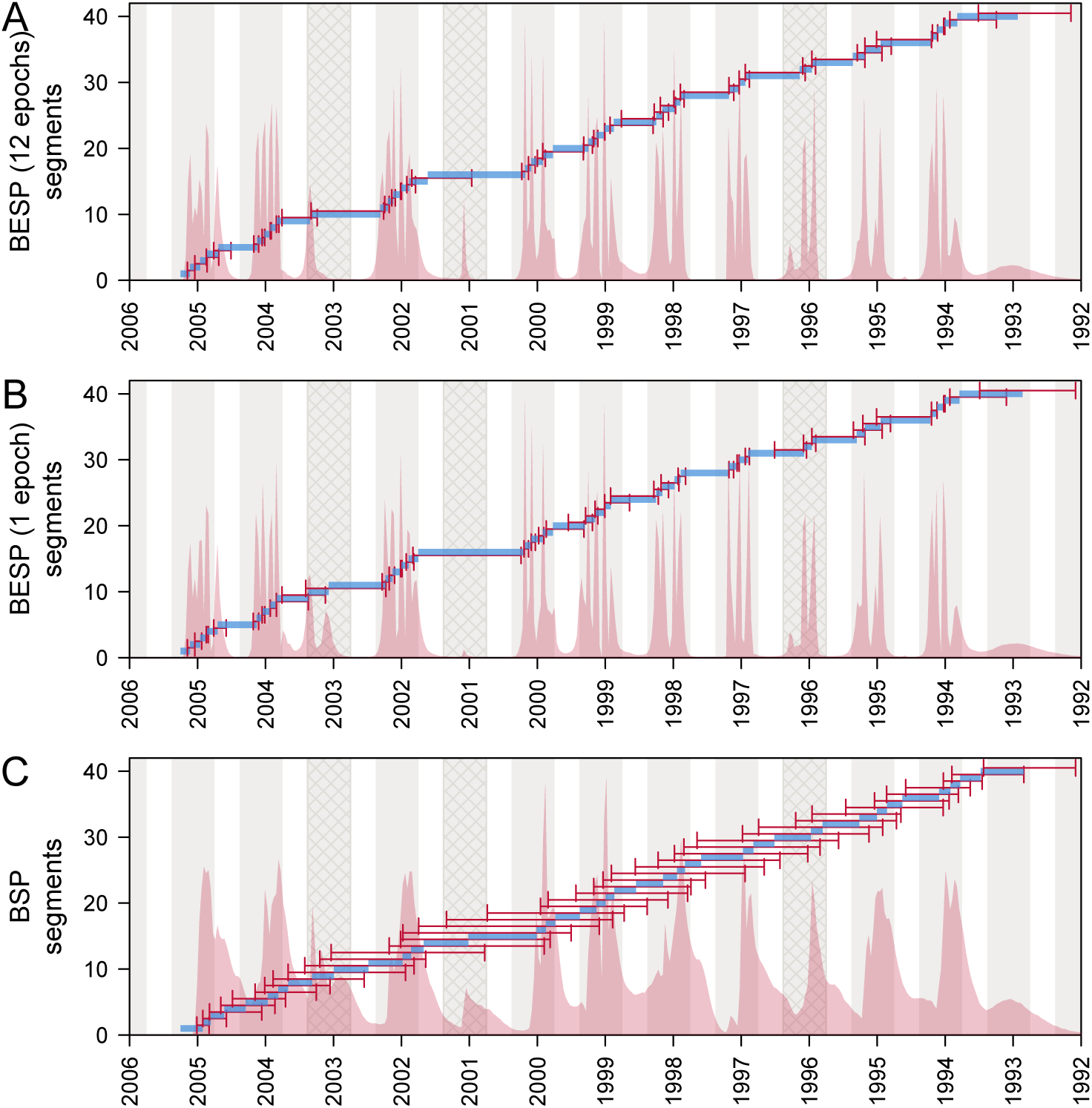
Population size segments for the alignment of 637 A/H3N2 HA sequences from NY state, as estimated under the 12-epoch BESP (A), single-epoch BESP (B) and BSP (C), with *p* = 40. Median posterior estimates of segments (*t*_*j*−1_–*t*_*j*_) are shown in blue. HPD intervals for the segment end-times are indicated by red arrows. Red shading shows the kernel density estimate of the posterior segment times (*t*_*j*_). Grey shading indicates the approximate period of influenza observation in New York state during each season (epidemiological week 40, to week 20 in the next year). Cross-hatched seasons are those where A/H3N2 was not the dominant influenza virus subtype.

**Fig. S8:**
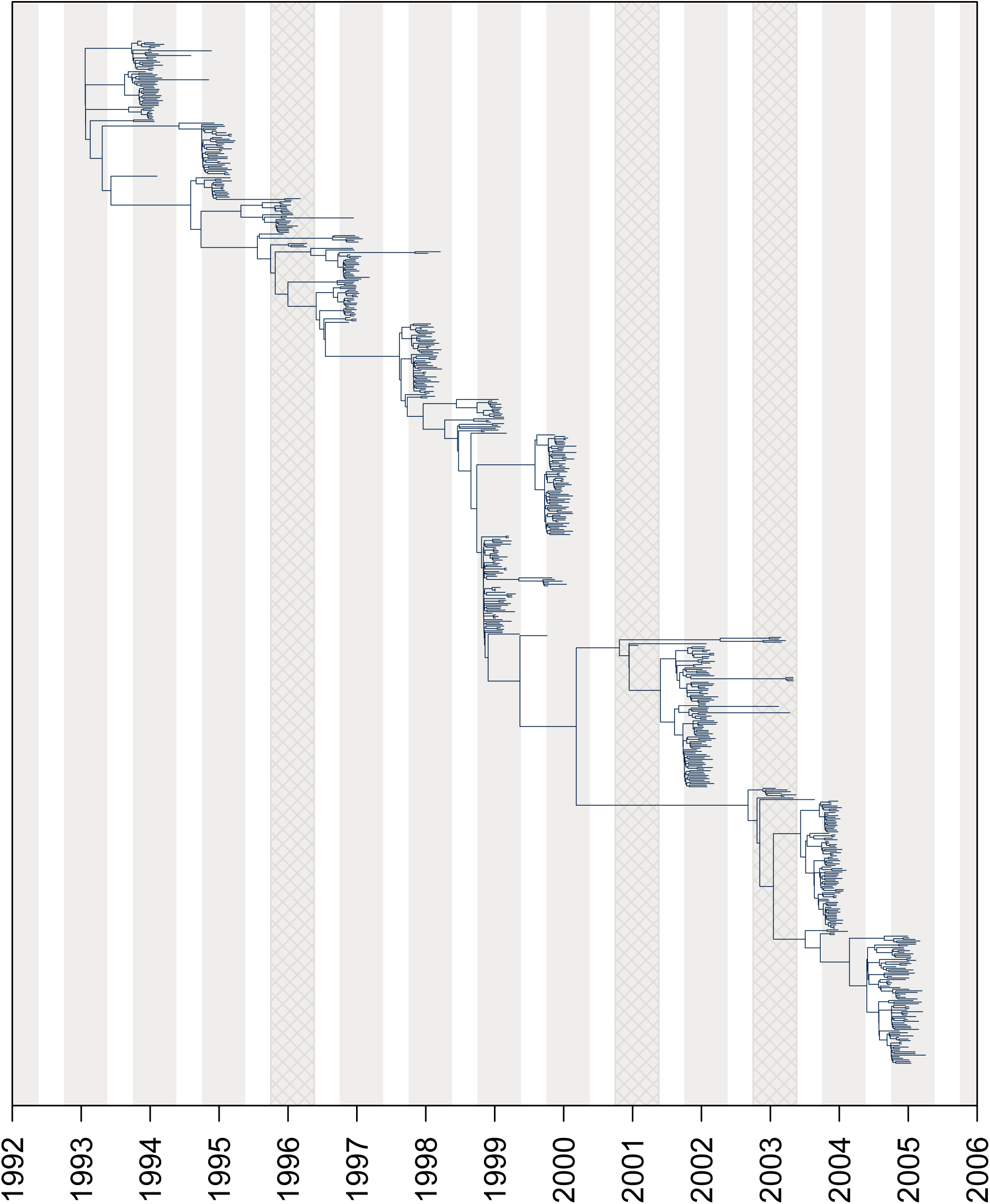
MCC tree of the alignment of 637 A/H3N2 HA sequences estimated under the 12-epoch BESP. Grey shading indicates the approximate period of influenza observation in New York state during each season (epidemiological week 40, to week 20 in the next year). Cross-hatched seasons are those where A/H3N2 was not the dominant influenza virus subtype.

**Fig. S9:**
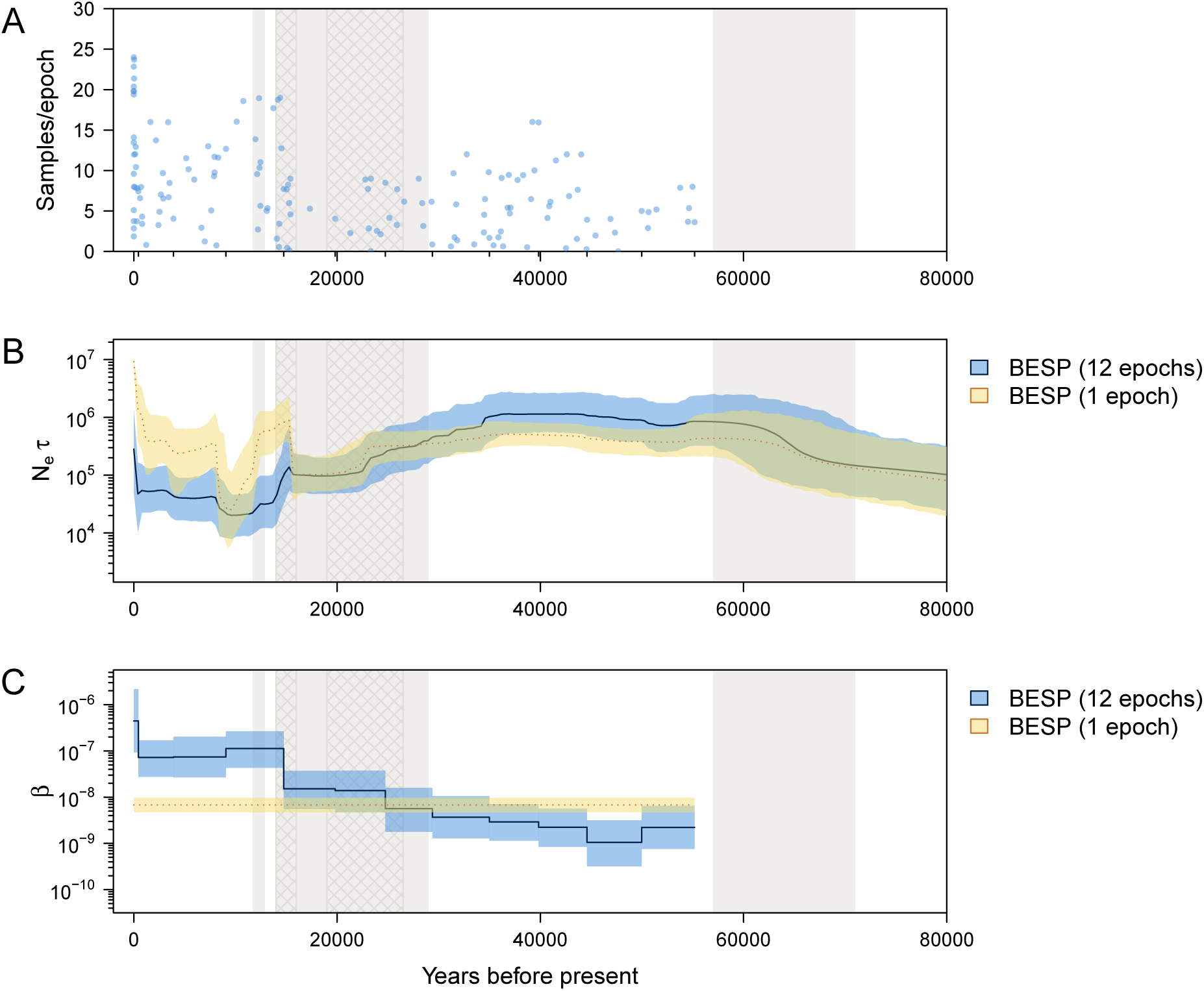
(A) Density of sequence sampling dates through time for the alignment of 152 bison mtDNA sequences that we used. Blue dots indicate stripcharts of individual samples for each sampling epoch. The height of the stripcharts is equal to the number of samples in each epoch. Small tick marks on the x-axis represent epoch times. Grey shading indicates cool periods in the Earth’s climate (from the present: Younger Dryas, Marine Isotope Stages (MIS) 2, MIS 4). The two cross hatched areas delimit the time of the last glacial maximum (≈26.5–19 ka BP) and approximate time of substantial human settlement of the Americas (≈16–14 ka BP). (B) Median (solid/dotted line) and 95% highest posterior density (HPD) intervals (shaded areas) for the genetic diversity estimates (*N*_*e*_*τ*) through time. The 12-epoch BESP estimate is shown in blue and the single-epoch BESP estimate in yellow. (C) Median (solid line/dotted line) and 95% HPD intervals (shaded areas) of the estimated sampling intensities (*β*) for each sampling epoch. The 12-epoch BESP estimates are in blue and a single-epoch (density-defined) estimate is in yellow.

**Fig. S10:**
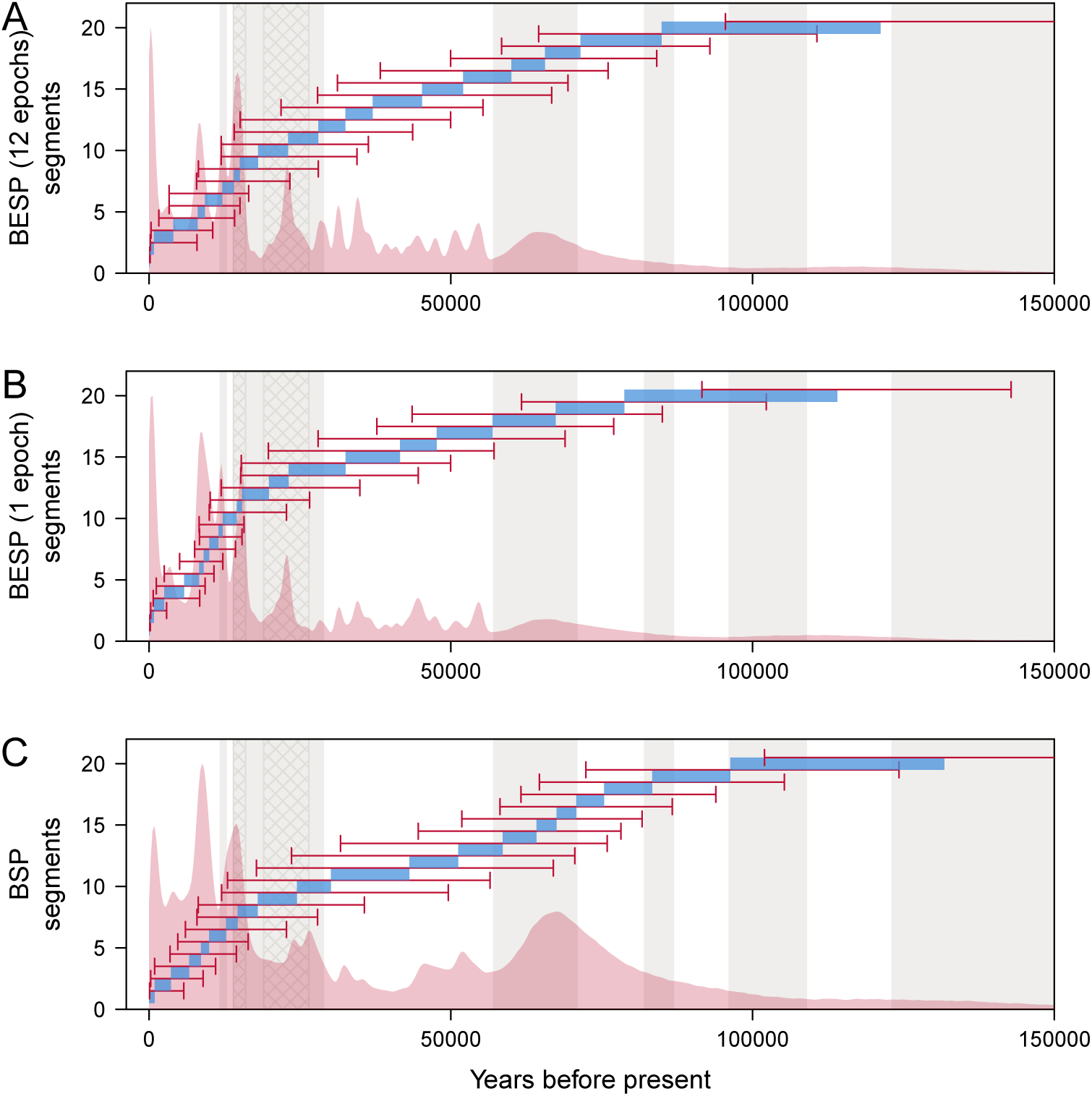
Population size segments for the alignment of 152 bison mtDNA sequences, as estimated under the 12-epoch BESP (A), single-epoch BESP (B) and BSP (C), with *p* = 20. Median posterior estimates of segments (*t*_*j*−1_–*t*_*j*_) are shown in blue. HPD intervals for the segment end-times are indicated by red arrows. Red shading shows the kernel density estimate of the posterior segment times (*t*_*j*_). Grey shading indicates cool periods in the Earth’s climate (from the present: Younger Dryas, Marine Isotope Stages (MIS) 2, MIS 4). The two cross hatched areas delimit the time of the last glacial maximum (≈26.5–19 ka BP) and approximate time of substantial human settlement of the Americas (≈16–14 ka BP).

**Fig. S11:**
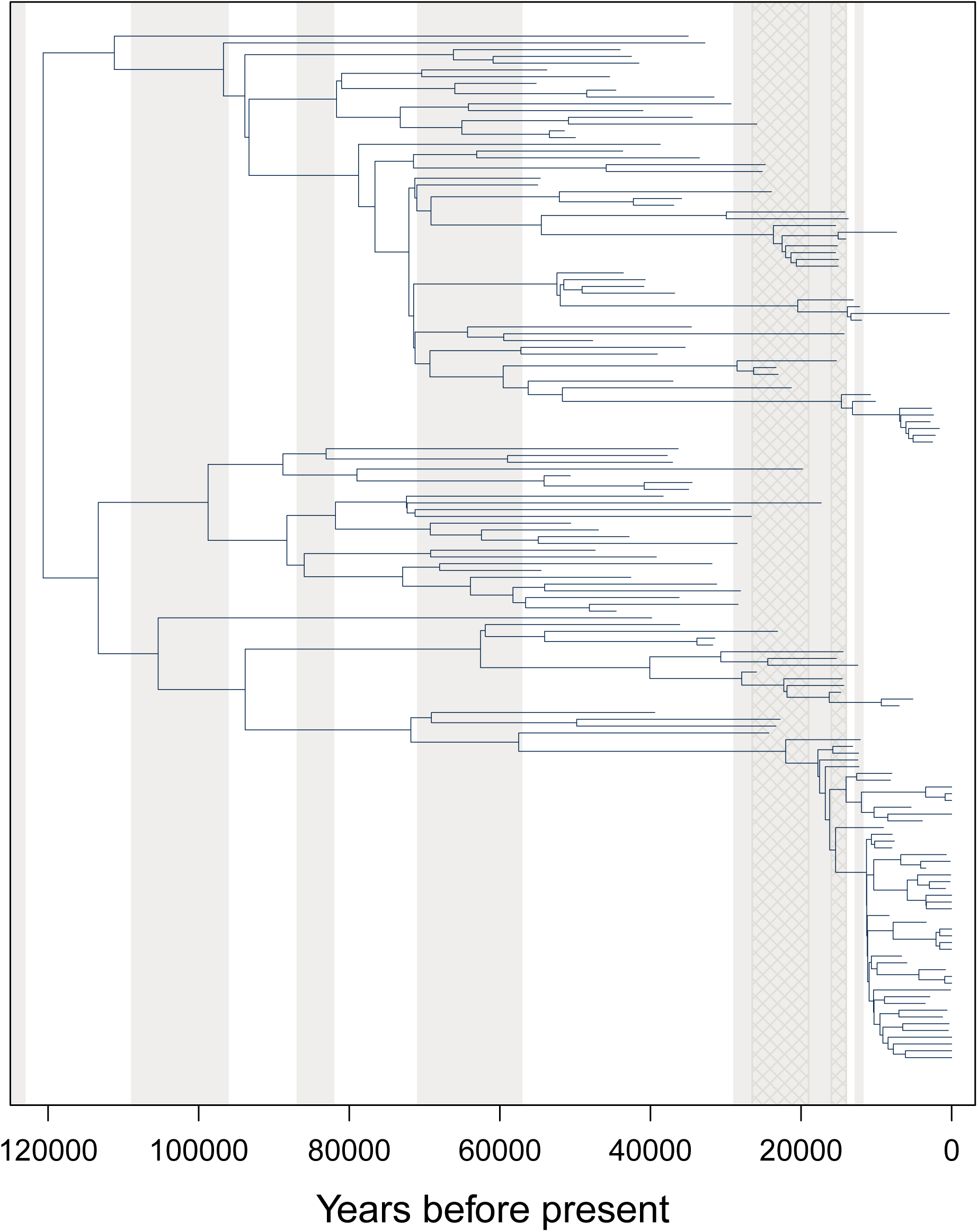
MCC tree of the alignment of 152 bison mtDNA sequences estimated under the 12-epoch BESP. Grey shading indicates cool periods in the Earth’s climate (from the present: Younger Dryas, Marine Isotope Stages (MIS) 2, MIS 4). The two cross hatched areas delimit the time of the last glacial maximum (≈26.5–19 ka BP) and approximate time of substantial human settlement of the Americas (≈16–14 ka BP).

## Footnotes

1 See https://github.com/laduplessis/BESP_paper-analyses/for simulated trees and inferred parameter trajectories for all replicates.

## References

Baele, G., Suchard, M., Rambaut, A., et al. 2017. Emerging Concepts of Data Integration in Pathogen Phylodynamics. Syst. Biol, 66(1): e47–65.

Biek, R., Pybus, O. G., Lloyd-Smith, J. O., and Didelot, X. 2015. Measurably evolving pathogens in the genomic era. Trends in Ecology & Evolution, 30(6): 306–313.

Bouckaert, R., Vaughan, T., Barido-Sottani, J., et al. 2019. BEAST 2.5: An Advanced Software Platform for Bayesian Evolutionary Analysis. PLoS Comp. Biol, 15(4): e1006650.

CDC 2019. Overview of influenza surveillance in the united states. https://www.cdc.gov/flu/weekly/overview.htm. [Online; last accessed 09-July-2019].

Drummond, A., Rambaut, A., Shapiro, B., et al. 2005. Bayesian Coalescent Inference of Past Population Dynamics from Molecular Sequences. Mol. Biol. Evol, 22(5): 1185–92.

Drummond, A. J., Pybus, O. G., Rambaut, A., Forsberg, R., and Rodrigo, A. G. 2003. Measurably evolving populations. Trends in Ecology & Evolution, 18(9): 481–488.

Drummond, A. J., Ho, S. Y. W., Phillips, M. J., and Rambaut, A. 2006. Relaxed Phylogenetics and Dating with Confidence. PLOS Biology, 4(5): e88.

Faulkner, J. R., Magee, A. F., Shapiro, B., and Minin, V. N. 2019. Horseshoe-based Bayesian nonparametric estimation of effective population size trajectories. Biometrics. In press.

Ferguson, N., Galvani, A., and Bush, R. 2003. Ecological and Immunological Determinants of Influenza Evolution. Nature, 422(6930): 428.

Gattepaille, L., Torsten, G., and Jakobsson, M. 2016. Inferring Past Effective Population Size from Distributions of Coalescent Times. Genetics, 204: 1191–206g.

Gill, M., Lemey, P., Faria, N., et al. 2012. Improving Bayesian Population Dynamics Inference: A Coalescent-Based Model for Multiple Loci. Mol. Biol. Evol, 30(3): 713–24.

Hall, M., Woolhouse, M., and Rambaut, A. 2016. The Effects of Sampling Strategy on the Quality of Reconstruction of Viral Population Dynamics using Bayesian Skyline Family Coalescent Methods: A Simulation Study. Virus Evol, 2(1).

Hasegawa, M., Kishino, H., and Yano, T.-a. 1985. Dating of the human-ape splitting by a molecular clock of mitochondrial DNA. Journal of Molecular Evolution, 22(2): 160–174.

Ho, S. and Shapiro, B. 2011. Skyline-plot Methods for Estimating Demographic History from Nucleotide Sequences. Mol. Ecol. Res, 11: 423–34.

Karcher, M., Palacios, J., Bedford, T., et al. 2016. Quantifying and Mitigating the Effect of Preferential Sampling on Phylodynamic Inference. PLoS Comp. Bio, 12(3).

Karcher, M., Palacios, J., Lan, S., et al. 2017. PHYLODYN: an R package for Phylodynamic Simulation and Inference. Mol. Ecol. Res, 17: 96–100.

Karcher, M., Suchard, M., Dudas, G., et al. 2019. Estimating Effective Population Size Changes from Preferentially Sampled Genetic Sequences. arXiv e-prints, page 1903.11797.

Kay, S. 1993. Fundamentals of Statistical Signal Processing: Estimation Theory. Prentice Hall.

Kingman, J. 1982. On the Genealogy of Large Populations. J. Appl. Prob, 19: 27–43.

Minin, V., Bloomquist, E., and Suchard, M. 2008. Smooth Skyride through a Rough Skyline: Bayesian Coalescent-Based Inference of Population Dynamics. Mol. Biol. Evol, 25(7): 1459–71.

Parag, K. and Pybus, O. 2017. Optimal Point Process Filtering and Estimation of the Coalescent Process. J. Theor. Biol, pages 153–67.

Parag, K. and Pybus, O. 2018. Exact Bayesian Inference for Phylogenetic Birth-death Models. Bioinformatics, 34(21): 3638–45.

Parag, K. and Pybus, O. 2019. Robust Design for Coalescent Model Inference. Syst. Biol, 68(5): 730–43.

Plummer, M., Best, N., Cowles, K., and Vines, K. 2006. Coda: Convergence diagnosis and output analysis for mcmc. R News, 6(1): 7–11.

Pybus, O. and Rambaut, A. 2009. Evolutionary Analysis of the Dynamics of Viral Infectious Disease. Nat. Rev Gen, 10: 240–50.

Pybus, O., Rambaut, A., and Harvey, P. 2000. An Integrated Framework for the Inference of Viral Population History from Reconstructed Genealogies. Genetics, 155: 1429–37.

Rambaut, A., Pybus, O., Nelson, M., et al. 2008. The Genomic and Epidemiological Dynamics of Human Influenza A Virus. Nature, 453(7195): 615–619.

Rambaut, A., Drummond, A., Xie, D., et al. 2018. Posterior Summarization in Bayesian Phylogenetics Using Tracer 1.7. Systematic Biology, 67(5): 901–904.

Rothenberg, T. 1971. Identification in Parametric Models. Econometrica, 39(3): 577–91.

Sagulenko, P., Puller, V., and Neher, R. A. 2018. TreeTime: Maximum-likelihood phylodynamic analysis. Virus Evolution, 4(1).

Shapiro, B. and Hofreiter, M. 2014. A Paleogenomic Perspective on Evolution and Gene Function: New Insights from Ancient DNA. Science, 343(6169): 1236573.

Shapiro, B., Drummond, A., Rambaut, A., et al. 2004. Rise and Fall of the Beringian Steppe Bison. Science, 306(5701): 1561–1565.

Shapiro, B., Rambaut, A., and Drummond, A. J. 2006. Choosing Appropriate Substitution Models for the Phylogenetic Analysis of Protein-Coding Sequences. Molecular Biology and Evolution, 23(1): 7–9.

Snyder, D. and Miller, M. 1991. Random Point Processes in Time and Space. Springer-Verlag, 2 edition.

Stack, J., Welch, J., Ferrari, M., et al. 2010. Protocols for Sampling Viral Sequences to Study Epidemic Dynamics. J. R. Soc. Interface, 7: 1119–27.

Stadler, T., Kuhnert, D., Bonhoeffer, S., et al. 2013. Birth-death Skyline Plot reveals Temporal Changes of Epidemic Spread in HIV and Hepatitis C Virus (hcv). PNAS, 110(1): 228–33.

Stamatakis, A. 2014. RAxML version 8: a tool for phylogenetic analysis and post–analysis of large phylogenies. Bioinformatics, 30(9): 1312–1313.

Strimmer, K. and Pybus, O. 2001. Exploring the Demographic History of DNA Sequences using the Generalized Skyline Plot. Mol. Biol. Evol, 18(12): 2298–305.

Viboud, C., Sun, K., Gaffey, R., et al. 2018. The RAPIDD Ebola Forecasting Challenge: Synthesis and Lessons Learnt. Epidemics, 22: 13–21.

Volz, E. and Frost, S. 2014. Sampling through Time and Phylodynamic Inference with Coalescent and Birth–death Models. J. R. Soc. Interface, 11(20140945).

WHO 2018. Fact sheet on seasonal influenza. https://www.who.int/en/news-room/fact-sheets/detail/influenza-(seasonal). [Online; last accessed 25-July-2019].

